# Bioinspired Silk Fibroin Mineralization for Advanced *In Vitro* Bone Remodeling Models

**DOI:** 10.1101/2022.06.17.496534

**Authors:** Bregje W.M de Wildt, Robin van der Meijden, Paul A.A. Bartels, Nico A.J.M. Sommerdijk, Anat Akiva, Keita Ito, Sandra Hofmann

## Abstract

Human *in vitro* bone models can create the possibility for investigation of physiological bone remodeling while addressing the principle of replacement, reduction and refinement of animal experiments (3R). Current *in vitro* models lack cell-matrix interactions and their spatiotemporal complexity. To facilitate these analyses, a bone-mimetic template was developed in this study, inspired by bone’s extracellular matrix composition and organization. Silk fibroin (SF) was used as an organic matrix, poly-aspartic acid (pAsp) was used to mimic the functionality of non-collagenous proteins, and 10x simulated body fluid served as mineralization solution. By using pAsp in the mineralization solution, minerals were guided towards the SF material resulting in mineralization inside and as a coating on top of the SF. After cytocompatibility testing, remodeling experiments were performed in which mineralized scaffold remodeling by osteoclasts and osteoblasts was tracked with non-destructive micro-computed tomography and medium analyses over a period of 42 days. The mineralized scaffolds supported osteoclastic resorption and osteoblastic mineralization, in the physiological bone remodeling specific sequence. This model could therefore facilitate the investigation of cell-matrix interactions and may thus reduce animal experiments and advance *in vitro* drug testing for bone remodeling pathologies like osteoporosis, where cell-matrix interactions need to be targeted.

## 1. Introduction

Bone is a highly dynamic tissue with multiple mechanical and metabolic functions that are maintained by the process of bone remodeling. Physiological bone remodeling follows a specific sequence of events: activation, bone resorption by osteoclasts, reversal, and bone formation by osteoblasts ^[1]^. Unbalanced bone remodeling can result in pathologies such as osteoporosis and osteopetrosis. Studies of these bone pathologies and their drug development are routinely performed in animal models. However, animal models represent human physiology insufficiently which is likely one of the reasons that only 9.6% of preclinically developed drugs are approved for regular clinical use ^[2,3]^. Human *in vitro* bone models can potentially facilitate the investigation of physiological human bone remodeling while addressing the principle of replacement, reduction, and refinement of animal experiments. Current studies aiming at mimicking bone remodeling mostly use osteoblast-osteoclast (progenitor) co-cultures to study indirect or direct cell-cell interactions in two dimensions (2D) ^[4–9]^. Although these studies have improved the understanding in factors involved in bone remodeling, they do not allow for studying the interactions with a three-dimensional (3D) complex bone-like matrix ^[10]^. Researchers that have attempted to mimic bone remodeling in 3D often 1) neglect the specific sequence of events *(i.e.* resorption, transition, formation (Figure 1A)) by starting their culture with osteoblast (progenitors) ^[11,12]^, or 2) only look at osteoclast and osteoblast markers with *e.g.* gene expression or enzymatic activity assays rather than at their function to resorb and form a bone-like matrix ^[13,14]^. As such, functional cell-matrix interactions and their temporal dynamics are often neglected ^[1]^ (Figure 1B). To enable the investigation of functional cell-matrix interactions and to mimic the sequence of these interactions *in vitro*, a bone-mimetic template is required ^[1,15–18]^.

**Figure 1.**
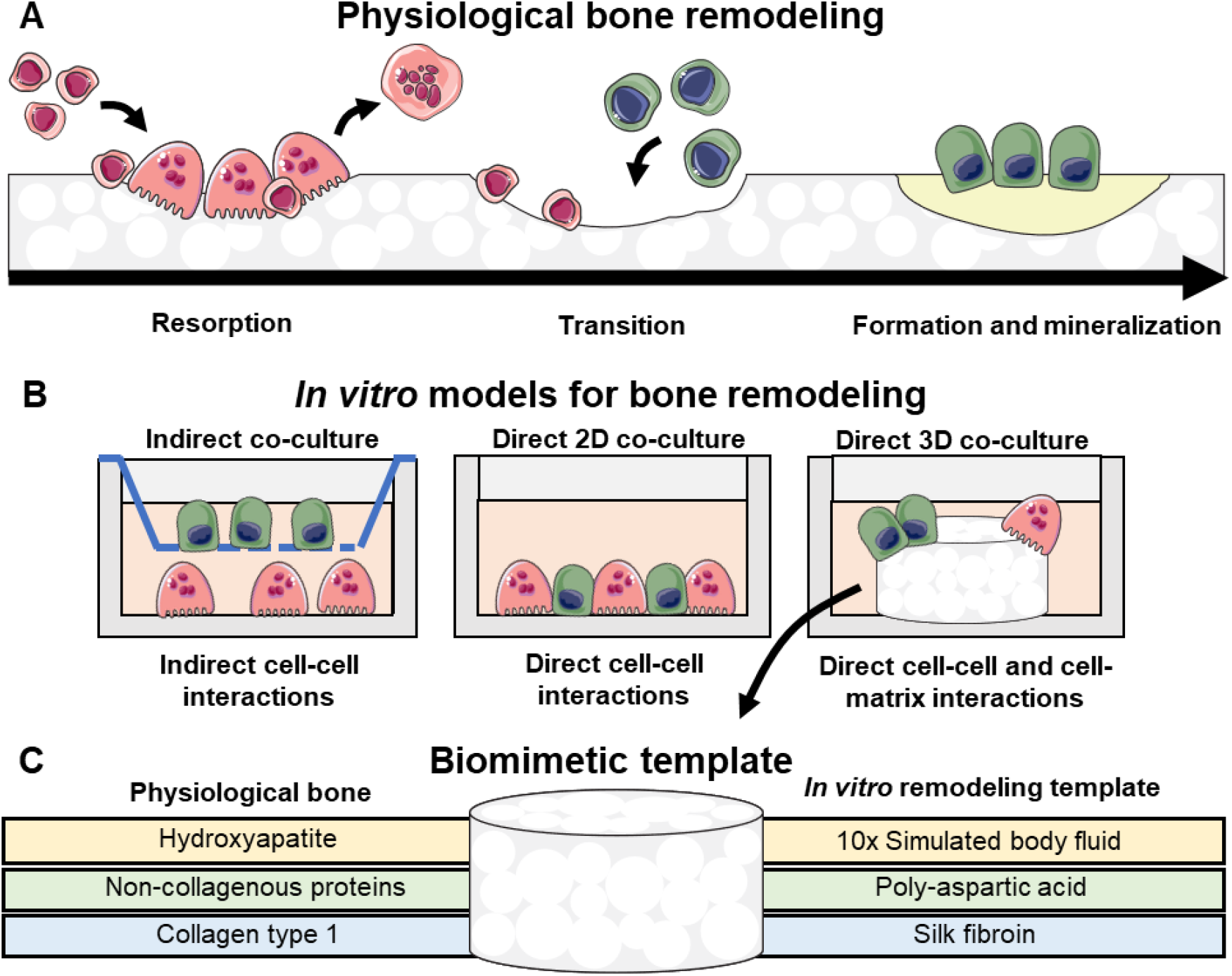
Reasoning towards the work presented in this study. (**A**) The physiological bone remodeling cycle starting with resorption after activation, then there is a transition phase followed by formation, mineralization and subsequent termination. (**B**) Current *in vitro* models for bone remodeling lack the investigation of cell-matrix interactions. (**C**) The proposed biomimetic template including the components present in physiological bone. Abbreviations: two-dimensional (2D), three-dimensional (3D). The figure was modified from Servier Medical Art, licensed under a Creative Common Attribution 3.0 Generic License (http://smart.servier.com/, accessed on 8 July 2021).

Bone tissue consists mainly of organic collagen type 1 and the inorganic mineral carbonated hydroxyapatite, which are highly organized at multiple hierarchical levels ^[19]^. Collagen mineralization starts when mineral precursors enter the collagen gap regions where carbonated hydroxyapatite crystals nucleate and grow outside the dimensions of the collagen fibril, resulting in mineralization inside (intrafibrillar) and outside (extrafibrillar) the collagen fibrils ^[20]^. A bone-mimetic template should include these characteristics. While the use of collagen type 1 as organic matrix seems obvious, drawbacks are the high biodegradability, low mechanical strength, and the difficulty of *in vitro* collagen self-assembly resulting in poorly organized low-density networks ^[21,22]^. The fibrous protein silk fibroin (SF) is a suitable organic alternative, thanks to its excellent mechanical properties, ease to process, and biocompatibility ^[23]^. SF features a unique structure which consists of hydrophobic *β*-sheets and hydrophilic amorphous acidic spacers, of which the latter could act as nucleation sites for mineral crystals similar to the collagen gap regions in bone ^[24]^. To mineralize SF, simulated body fluid (SBF) has been widely used ^[25]^. Immersing materials in this solution containing physiological ion concentrations results in the formation of calcium phosphate crystalline structures like apatite found in real bone ^[26]^. However, material mineralization with SBF could take up to 4 weeks and requires frequent replenishment of the solution ^[26,27]^. This mineralization period often only results in a non-uniform mineral coating, rather than infiltration of minerals into the material’s structure ^[25]^. *In vivo*, non-collagenous proteins are believed to play an instrumental role in the infiltration of mineral precursors into collagen fibrils ^[28]^. In bone tissue, extracellular levels of calcium and phosphate ions are supersaturated and their precipitation is therefore controlled by these acidic proteins ^[28]^. *In vitro*, poly-aspartic acid (pAsp) can be used to mimic the functionality of these acidic non-collagenous proteins as its addition to a mineralization solution has been shown to induce intrafibrillar collagen mineralization ^[29]^. As such, pAsp might improve mineral distribution and infiltration for SF as well. Therefore, in this study a bone-mimetic template was developed using SF as organic material mineralized with SBF under influence of pAsp (Figure 1C).

To accomplish this, we evaluated the use of pAsp as a substitute to the mineralization solution and/or integrated into the SF material to improve SF mineralization. The effects of these material preparation methods on material cytocompatibility were subsequently tested in two-dimensional (2D) films for human monocytes (MCs) and mesenchymal stromal cells (MSCs) as the osteoclast and osteoblast progenitors, respectively. Improved mineralization methods were also applied to and evaluated in three-dimensional (3D) porous SF scaffolds. *In vitro* remodeling of these scaffolds by human osteoclastogenically stimulated MCs and osteogenically stimulated MSCs was tracked longitudinally which enabled the investigation of cell-matrix interactions and their temporal dynamics. As a result, pAsp was instrumental for SF mineralization in a similar manner as for collagen mineralization. Mineralized SF scaffolds supported osteoclastic resorption and enhanced osteoblastic mineralization. As such, our model allowed for investigating functional cell-matrix interactions and their dynamics and may therefore advance *in vitro* drug testing for bone remodeling pathologies like osteoporosis, where cell-matrix interactions need to be targeted.

## 2. Results

### 2.1 Mineralization optimization and characterization of silk fibroin films

While intrafibrillar mineralization of small amounts of collagen using pAsp as a nucleation inhibiter in the mineralization solution has been established, large scale intrafibrillar mineralization of collagen scaffolds is still challenging ^[30,31]^. The use of nucleation inhibitors only in solution does not fully represent the physiological situation in which non-collagen proteins are bound to the matrix and might thus not provide the optimal conditions for homogeneous scaffold mineralization ^[32]^. Therefore, we choose to not only study the effect of pAsp in the mineralization solution on SF mineralization, but we also mixed it into the SF (Figure 2A). To enable the screening of multiple parameters and to facilitate the analyses, mineralization optimization and characterization was performed in 2D. Pure SF (SF w/o pAsp) and SF containing 5 wt% pAsp (SF w/5% pAsp) solutions were casted to form films with a diameter of 10 mm and a thickness of ~300 μm. To check for the presence of pAsp in SF w/5% pAsp films, films were stained with the cationic dye alcian blue to allow for visualization of the negatively charged pAsp. The addition of pAsp indeed led to a more intense blue stain in SF films with 5% pAsp when compared to plain SF films (Figure S1). The presence of a small amount of pAsp in the SF material was also confirmed by chemical analyses. Raman spectroscopy measurements revealed a small peak at 1783 cm^-1^, suggesting the presence of pAsp (Figure S2). X-ray photoelectron spectroscopy (XPS) measurements revealed a carbon peak with wider shape, which is likely attributed to the carboxyl group in pAsp (Figure S3). By measuring the water contact angle an increase in hydrophilicity of SF w/5% pAsp was observed by a decrease in water contact angle (Figure 2B). Both types of films were subsequently mineralized using 10x SBF ^[27]^ (SBF) or 10x SBF with 100 μg/ml pAsp (SBF-pAsp). Films were mineralized in this solution for either one week (W1, no replenishment of mineralization solution) or two weeks (W2, one mid-way replenishment of mineralization solution). Baseline films were used as non-mineralized controls (NM-control) (Figure 2A). After mineralization, the property of pAsp to prevent mineral precipitation in solution was verified by measuring the optical density of the mineralization media. The addition of pAsp to the mineralization solution indeed led to a statistically significant decrease in mineralization solution optical density (Figure 2C). Mineralization solution optical density was also significantly decreased after one mineralization solution replenishment (W2). In the films where pAsp was added to the mineralization solution the optical density after W2 reduced towards almost the optical density of ultra-pure water (UPW). Most likely, optical density was reduced after W2 because some calcium phosphate crystals were already nucleated on the film to which calcium and phosphate ions could precipitate more easily ^[33]^. A reverse effect was found for the calcium content (Figure 2D). Both the addition of pAsp to the mineralization solution as well as the replenishment of the solution resulted in a statistically significant increase in calcium content of the film, whereas the addition of pAsp to the SF material did not affect its mineralization. Calcium content results were confirmed by alizarin red staining of film cross-sections with a clear red staining on top of films mineralized with pAsp in the mineralization solution after W2 (Figure S4E+J, cross-sections). In these groups, only mineralized SF w/o pAsp films showed red staining inside the film indicating mineral infiltration into the films (Figure S4E, cross-section). While the addition of pAsp to the material did not affect its mineralization, it caused a statistically significant decrease in Young’s modulus (stiffness) compared to plain SF films, as measured with nanoindentation (Figure 2E).

**Figure 2.**
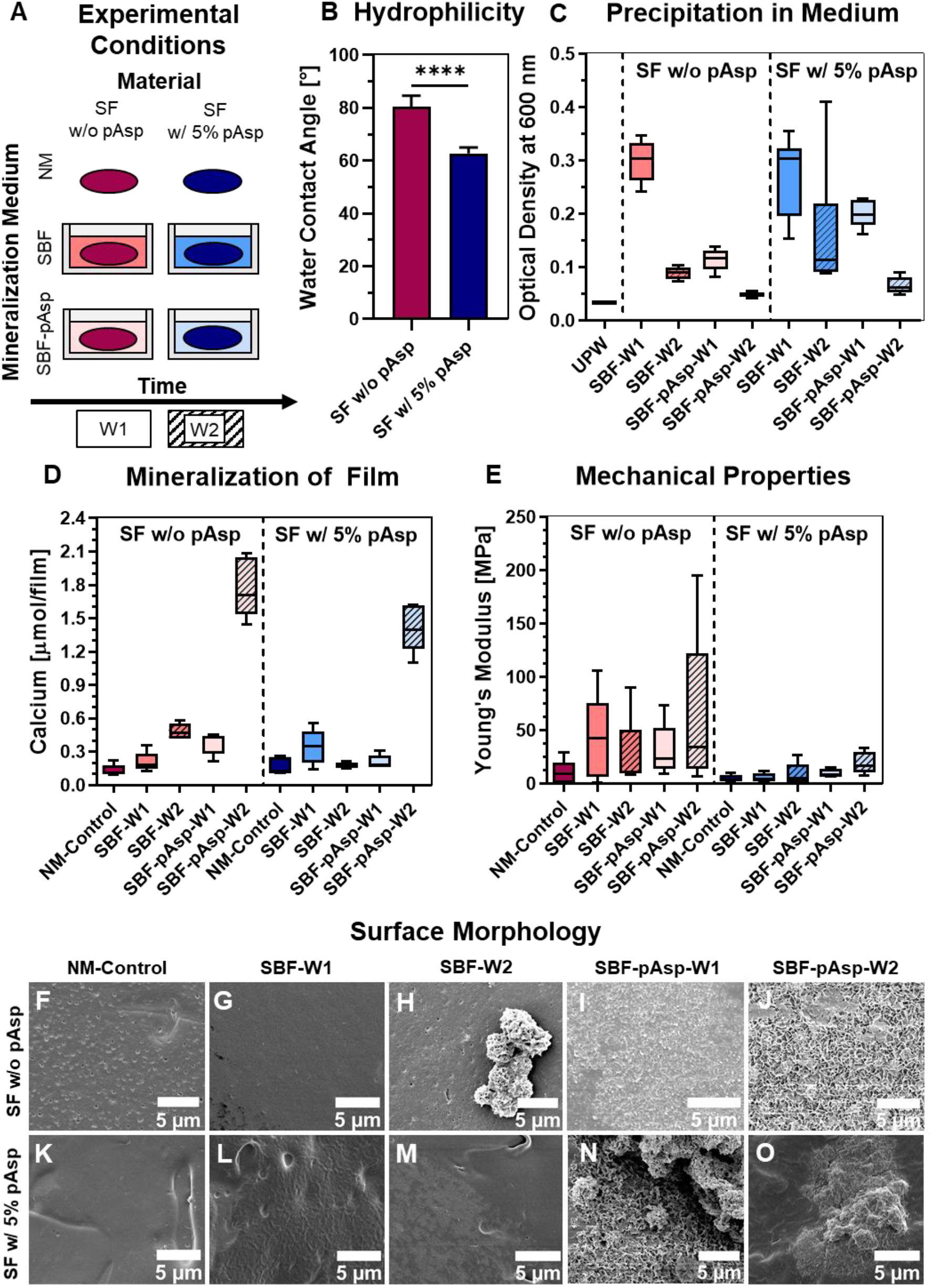
Mineralization optimization and characterization of SF films. (**A**) Experimental variables included in the mineralization optimization. (**B**) Water contact angle quantification, *p*<0.0001 (Independent t-test). (**C**) Solution optical density measurement to detect mineral precipitation, *p*<0.0001 for mineralization time and mineralization solution (Kruskal-Wallis test for main effects with Bonferroni correction for multiple comparisons). (**D**) Mineral quantification in film, measured by calcium content, *p*<0.01 for mineralization time and *p*<0.05 for mineralization solution (Kruskal-Wallis test for main effects with Bonferroni correction for multiple comparisons). (**E**) Stiffness measured with nanoindentation, *p*<0.01 for the material (Kruskal-Wallis test for main effects with Bonferroni correction for multiple comparisons) (**F-O**) Surface and mineral morphology visualized with scanning electron microscopy. Abbreviations: silk fibroin (SF), simulated body fluid (SBF), poly-aspartic acid (pAsp), non-mineralized (NM), week (W), ultra-pure water (UPW).

When visualizing the film surfaces with scanning electron microscopy (SEM), non-mineralized SF w/5% pAsp films had a rougher surface than SF w/o pAsp films (Figure 2F+K). Mineral crystals on the surface were observed in SF w/o pAsp films mineralized with only SBF after W2, and in all films mineralized with pAsp in the solution after W1 and W2 (Figure 2H-J and 2N-O). As mineralization duration (W2) and the addition of pAsp to the solution positively influenced mineralization of the films, these mineralization conditions were used for 3D scaffold mineralization and cytocompatibility testing of the 2D films. Although the addition of pAsp to the material did not improve mineralization, this condition was still included to investigate the influence of bound pAsp on mineral distribution throughout the scaffold. In addition, the increased hydrophilicity and roughness of the SF w/5% pAsp films might still be beneficial for cell proliferation, osteoprogenitor differentiation, and osteoclastic resorption ^[34–37]^.

### 2.2 Characterization of mineralized silk fibroin scaffolds

SF scaffolds w/o pAsp and SF w/5% pAsp were mineralized using a mineralization solution of SBF with 100 μg/ml pAsp for 2 weeks with one solution replenishment after one week. Like in the films, calcium was detected in the mineralized scaffolds with no differences between the SF w/o pAsp and SF w/5% pAsp (Figure 3A). Mineralization led to an increased scaffold stiffness measured with an unconfined compression test (Figure 3B). Although not statistically significant, the addition of pAsp to the material seemed to negatively influence the average stiffness, something that was also observed in the films. Next, the scaffolds were analyzed for mineral distribution. Because of the radiolucent nature of SF when immersed in water, mineralization could be localized with micro-computed tomography (*μ*CT) scanning of the scaffolds. It was hypothesized that the addition of pAsp to the material could lead to improved mineral distribution throughout the scaffold. However, a positive influence of the addition of pAsp to the scaffold on mineral distribution could not be detected (Figure 3C and Figure S5). In both the radiographs and the quantification of the percentage minerals present in the central ~8% scaffold volume, no clear differences were found between the two material types.

**Figure 3.**
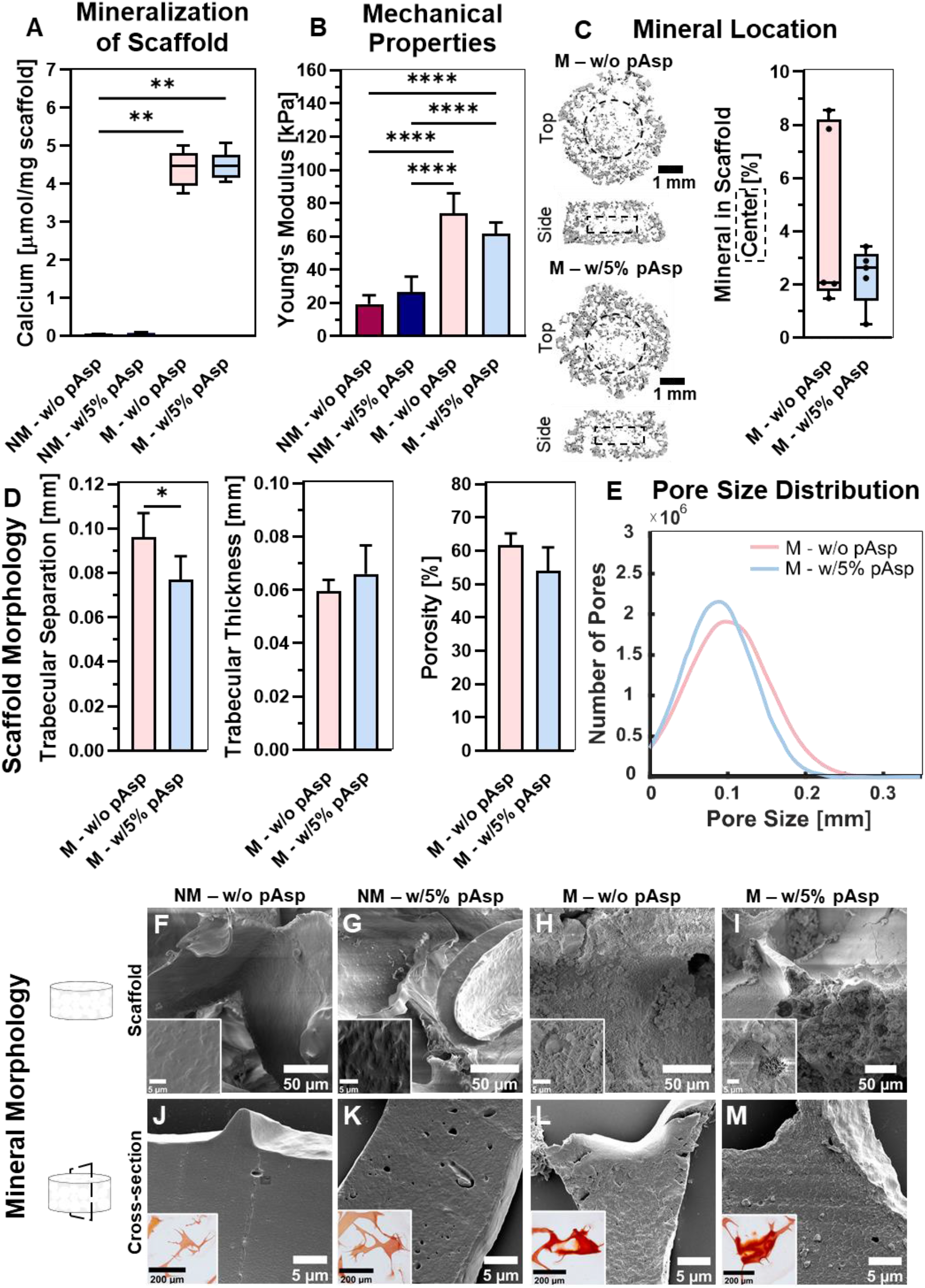
Characterization of mineralized SF scaffolds. (**A**) Mineral quantification in scaffold, measured by calcium content, *p*<0.05 (Kruskal-Wallis and Dunn’s post hoc tests). (**B**) Stiffness measured with a full unconfined compression test, *p*<0.05 (One-way ANOVA and Holm-Šídák’s post hoc tests). (**C**) Mineral location visualized with *μ*CT and quantified in the central ~8% scaffold volume to indicate mineral distribution. Dashed boxes represent the segmented part for mineral quantification in the center, *ns* (Mann-Whitney U). (**D**) Quantified scaffold morphology obtained with *μ*CT, including the trabecular separation (average pore size), *p*<0.05 (Independent t-test), trabecular thickness, *ns* (Independent t-test), porosity, *ns* (Independent t-test), and (**E**) the pore size distribution (gaussian fit). (**F-I**) Mineral morphology on scaffold surface visualized with SEM. (**J-M**) Morphology visualized with SEM and micrographs of calcium localization (insert, alizarin red staining) of cross-sections from scaffolds. (\**p*<0.05, \*\**p*<0.01, \*\*\**p*<0.001, \*\*\*\**p*<0.0001). Abbreviations: poly-aspartic acid (pAsp), non-mineralized (NM), mineralized (M), silk fibroin (SF), micro-computed tomography (*μ*CT), scanning electron microscopy (SEM).

By drying the mineralized scaffolds, their 3D morphology could be characterized after *μ*CT scanning (Figure 3D and Figure S5). These analyses revealed a smaller average pore size per scaffold, measured as trabecular separation. In the distribution of individual pore diameters, no clear differences were observed (Figure 3E). Although not significant, the decrease in average trabecular separation by the addition of pAsp to the material seemed reflected by an increase in trabecular thickness, trabecular number and trabecular connectivity density, and a decrease in porosity (Figure 3D and Figure S5). We then studied the mineral morphology with SEM. Minerals in the plain SF scaffolds (Figure 3H) appeared more homogeneously distributed in a layer on the surface when compared to minerals in SF w/5% pAsp scaffolds (Figure 3I). On SF w/5% pAsp scaffolds, minerals appeared more often as chunks, which likely caused the differences in scaffold morphology parameters. Mineral infiltration seemed present in both scaffolds w/o pAsp and w/5% pAsp detected by alizarin red stained scaffold sections and a change in cross-section structure after mineralization (Figure 3L+M). To further investigate mineral infiltration into the SF material, Raman microscopy and spectroscopy were performed on scaffold cross-sections. The infiltration of mineral was observed in both SF w/o pAsp and SF w/5% pAsp scaffolds (Figure 4). Hydroxyapatite was observed throughout the whole scaffold trabecula, with a higher degree of mineralization at the surface of the trabecula indicated by the differences in the 960, 420 and 590 cm^-1^ areas representing the v_1_, v_2_ and v_4_ vibrations of hydroxyapatite respectively. In SF w/5% pAsp scaffolds, more mineral precipitation was observed at the trabecula surface. These minerals precipitated in the presence of pAsp, identified through the presence of the 1783 cm^-1^ peak (Figure 4D). XPS measurements revealed the presence of calcium, phosphate and pAsp in both mineralized SF w/o pAsp and SF w/5% pAsp scaffolds (Figure S3). The presence of pAsp was observed by the carbon peak with wider shape relative to non-mineralized SF w/o pAsp scaffolds indicative for the presence of the carboxyl group of pAsp. As such, pAsp was likely instrumental for the mineralization of both SF w/o pAsp and SF w/5% pAsp scaffolds.

**Figure 4.**
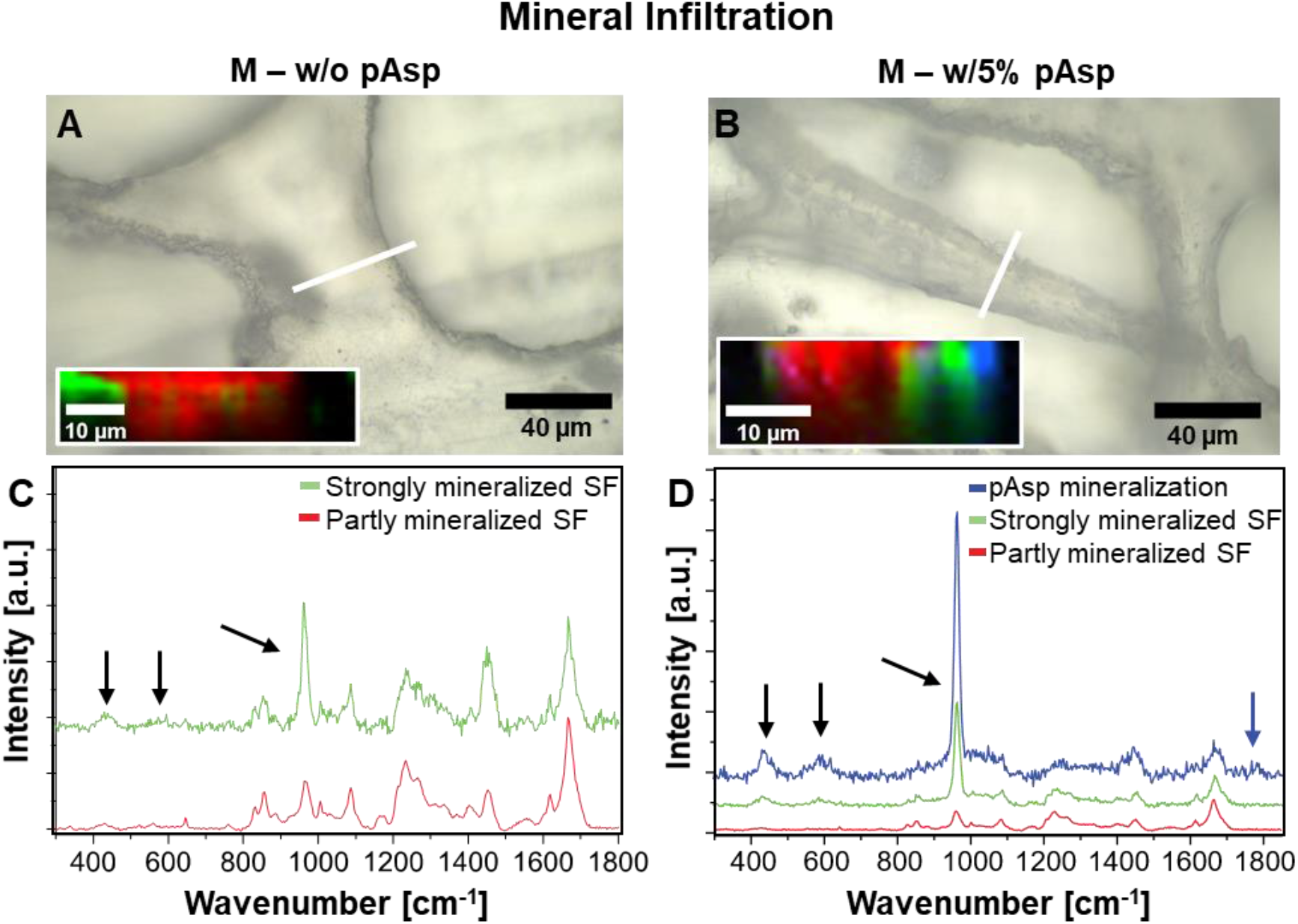
Raman microscopic analysis of mineralized SF scaffold sections to detect mineral infiltration. (**A**) Optical image of section of mineralized plain SF scaffold, scanned area highlighted in white. Insert in optical image presents the distribution of the strongly mineralized SF (green) and partly mineralized SF (red) in the analysed (50×12 μm) area. (**B**) Section of mineralized SF scaffold with 5% pAsp in the SF material. Insert in optical image presents the distribution of mineralized pAsp (blue), strongly mineralized SF (green) and partly mineralized SF (red). (**C**) Raman spectra of the mineralized plain SF scaffold. (**D**) Raman spectra of the mineralized SF scaffold with 5% pAsp added to the SF material. Mineralization with pAsp was identified through the presence of the 1783 cm^-1^ peak (blue arrow). (**C-D**) Black arrows indicate the 960, 420 and 590 cm^-1^ areas representing the v_1_, v_2_, v_4_, vibrations of hydroxyapatite. Abbreviations: poly-aspartic acid (pAsp), mineralized (M), silk fibroin (SF).

### 2.3 Cytocompatibility testing of mineralized silk fibroin films

Before the mineralized scaffolds were used for a co-culture experiment to study their remodeling *in vitro*, we first tested the materials’ cytocompatibility by running mono-cultures of human MCs and MSCs as the osteoclast and osteoblast progenitors, respectively. SF w/o pAsp and SF w/5% pAsp films, mineralized and non-mineralized, were evaluated for their cytocompatibility. For MSCs, the presence of w/5% pAsp in SF films seemed to negatively influence cell content which could be observed from DNA content measurements after 7 days culture (Figure 5C). These results were confirmed by micrographs of nuclei and F-actin staining from day 7, with clearly most cells present on mineralized SF w/o pAsp films (Figure 5D-G). To check whether these observations are a result of proliferation, cell death, or cell attachment, metabolic activity and cytotoxicity *(i.e.* cell death) were tracked over time. For the metabolic activity, the conversion of resazurin to fluorescent resorufin by viable cells was measured. These results reflected the DNA measurements, with highest metabolic activity in SF w/o pAsp films over the entire culture period (Figure 5A). Supernatant lactate dehydrogenase (LDH) activity, which is an intracellular enzyme released into the medium upon cell death, did not reveal clear cytotoxic effects of the different films (Figure 5B). Differences in cytotoxicity could be explained by the number of cells present on these films as indicated above. This indicates that the higher number of cells on SF w/o pAsp films is the result of cell attachment rather than more proliferation on these films or more cell death in SF w/5% pAsp films. This is in contrast with literature reporting often a positive influence of hydrophilicity on cell attachment ^[38,39]^. Interestingly, on mineralized SF w/o pAsp films, small particles in the proximity of cells were observed with SEM, which might indicate the presence of mineral nodules or matrix vesicles ^[40]^ (Figure 5J, white arrows). For MCs, no clear effects of the addition of pAsp to the material nor of mineralization were found in terms of DNA content on day 7 (Figure 5N). These results were in line with the metabolic activity measurements and with micrographs of a nuclei and F-actin staining from day 7 (Figure 5L and 5O-R). Only cytotoxicity in MCs cultured on mineralized SF w/5% pAsp films seemed higher than in MCs cultured on a film where pAsp was not added to the material (Figure 5M). This effect was however only observed on day 2 and day 7. After a period of 7 days, multinucleated osteoclast-like cells were observed in all conditions (Figure 5O-R, white arrows). To check whether these osteoclast-like cells also had the capability to resorb the material, films were visualized with SEM. In mineralized films, resorption pits were indeed observed indicating osteoclastic resorption (Figure 5U+V, white arrows). Resorption pits seemed largest in SF w/5% pAsp films (Figure 5V), indicating that resorption might be enhanced by the increased hydrophilicity as earlier observed ^[36]^. Based on these cytocompatibility evaluations, both SF w/o pAsp and SF w/5% pAsp can be considered suitable for cell culture with human MCs and MSCs. Because the addition of pAsp to the material led to reduced MSC attachment, a decreased pore size, and a heterogeneous mineral morphology, pAsp was left out the material for the 3D *in vitro* remodeling model.

**Figure 5.**
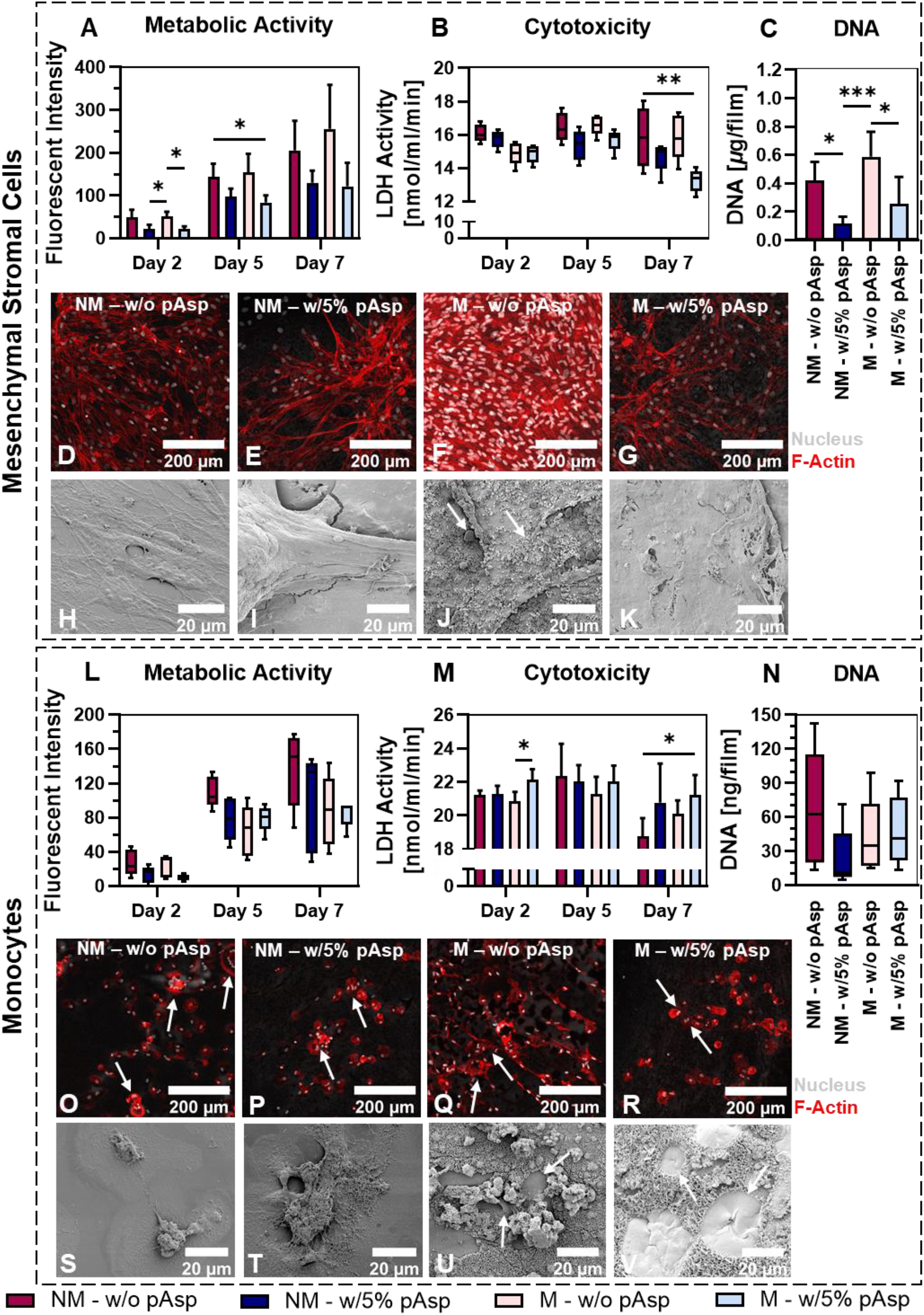
Cytocompatibility testing of mineralized SF films. (**A**) Metabolic activity measurements of MSCs using PrestoBlue™, *p*<0.05 (Two-way ANOVA and Turkey’s post hoc tests within each time point). (**B**) Cytotoxicity (cell death) for MSCs measured by LDH release in the medium, *p*<0.05 (Kruskal-Wallis and Dunn’s post hoc tests). (**C**) DNA content per film for MSCs, *p*<0.05 (One-way ANOVA and Holm-Šídák’s post hoc tests). (**D-G**) Micrographs of MSCs stimulated to undergo osteogenic differentiation, stained for F-Actin (red) and the nucleus (gray). (**H-K**) Visualization of MSC layer on films with SEM. (**L**) Metabolic activity measurements of MCs, *ns* (Kruskal-Wallis test). (**M**) Cytotoxicity for MCs, *p*<0.05 (Two-way ANOVA and Turkey’s post hoc tests within each time point). (**N**) DNA content per film for MCs, *ns* (Kruskal-Wallis test). (**O-R**) Micrographs of MCs stimulated to undergo osteoclastic differentiation, stained for F-Actin (red) and the nucleus (gray). (**S-V**) Visualization of cells and resorption on films with SEM. (\**p*<0.05, \*\**p*<0.01, \*\*\**p*<0.001, \*\*\*\**p*<0.0001). Abbreviations: poly-aspartic acid (pAsp), non-mineralized (NM), mineralized (M), mesenchymal stromal cells (MSCs), monocytes (MCs), lactate dehydrogenase (LDH), scanning electron microscopy (SEM).

### 2.4 *In vitro* remodeling of mineralized silk fibroin scaffolds

To investigate whether our bioinspired mineralized SF scaffold could enable the *in vitro* investigation of cell-matrix interactions and their temporal dynamics as described for physiological bone remodeling, we performed a MC-MSC co-culture for 42 days (Figure 6A). On day 21, medium was switched from osteoclastogenic to osteogenic. To track the remodeling dynamics, constructs were weekly scanned with *μ*CT, cell supernatants were collected, and constructs were sacrificed for analyses at day 21 and 42. Cell supernatants or cell lysates were analyzed for resorption (tartrate resistant acid phosphatase (TRAP) ^[14,41]^), transition (LDH to indicate potential osteoclast apoptosis), and formation (alkaline phosphatase (ALP) and pro-collagen 1 c-terminal propeptide (PICP) ^[14]^) markers. First, the influence of *μ*CT scanning on cell death was evaluated over 21 days. No differences between scanned and unscanned constructs were found, *μ*CT scanning was therefore considered as a harmless method to track *in vitro* remodeling (Figure S6). From day 7 to day 28, elevated TRAP activity was measured whereafter TRAP activity reduced to baseline levels (Figure 6B). During this resorption phase, TRAP activity in mineralized co-cultured constructs was significantly higher from day 14 on. This indicates that the 3D mineralized surface promotes osteoclast activity, which was observed earlier ^[11]^. In addition to the TRAP measurements, mineral resorption, which could only be studied in radiopaque mineralized constructs, seemed increased during the same period and resorption sites were identified (Figure 6E+H, yellow arrows, and Figure S6).

**Figure 6.**
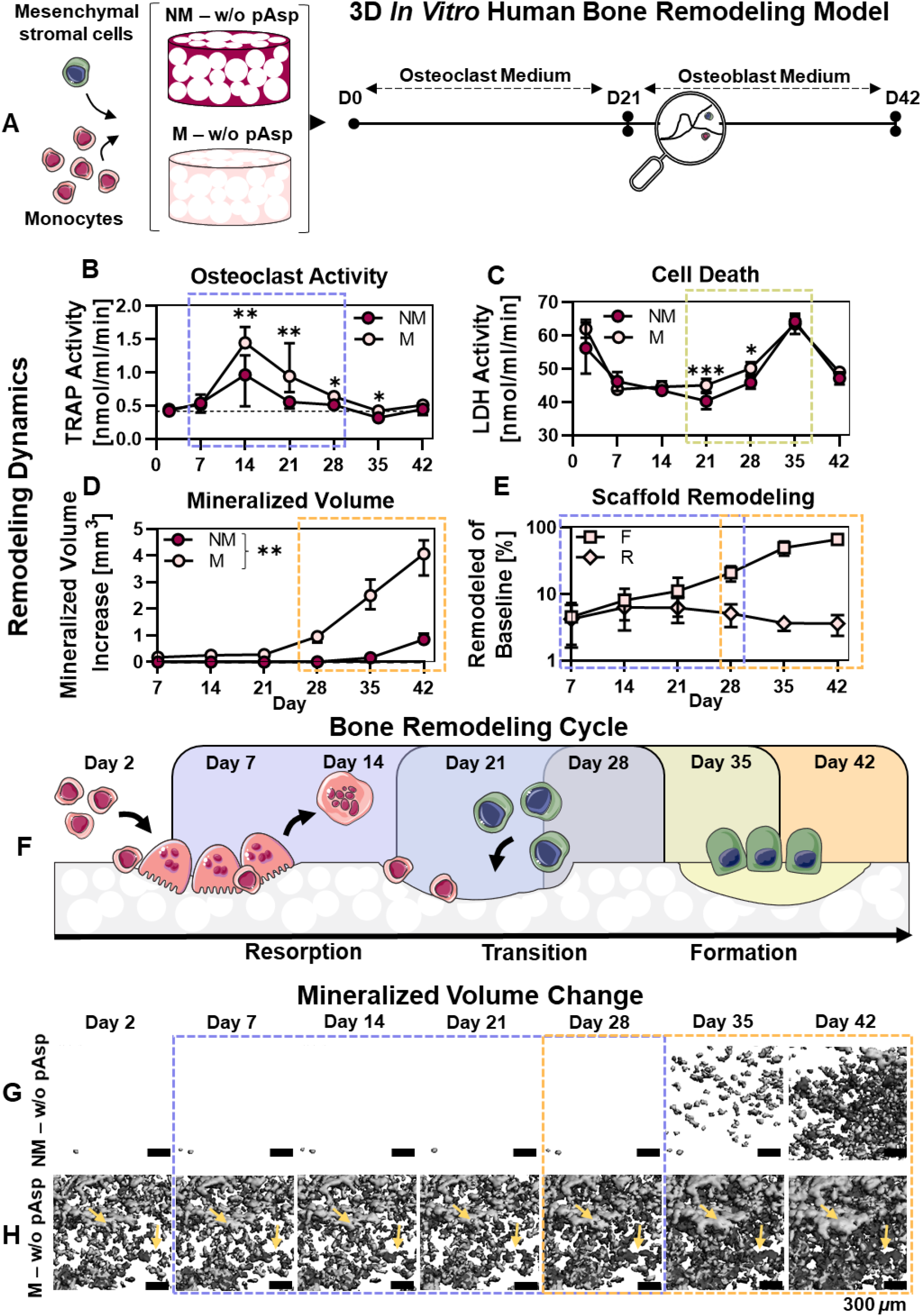
*In vitro* remodeling of mineralized silk fibroin scaffolds. (**A**) Experimental set-up. (**B**) TRAP activity quantification as a measure for osteoclast activity, dashed line represents the median value at baseline (day 2), *p*<0.05 (Mann-Whitney U tests per time point with Bonferroni correction for multiple comparisons). (**C**) Cell death measured by LDH release in the medium, *p*<0.05 (Two-way ANOVA and Turkey’s post hoc tests within each time point). (**D**) Mineralized volume measured with *μ*CT, *p*<0.01 for each time point (Mann-Whitney U tests per time point with Bonferroni correction for multiple comparisons). (**E**) Cumulative mineral formation (F) and resorption (R) as a percentage of the baseline scaffold, obtained after registration of *μ*CT scans. (**F**) The physiological bone remodeling cycle described in literature and the similarities to the remodeling dynamics found in the presented model. (**G**) Mineralization over time visualized with *μ*CT for non-mineralized scaffolds. (**H**) Resorption and mineralization over time visualized with *μ*CT for mineralized scaffolds. Yellow arrows represent remodeling/coupling sites. Dashed boxes in figure represent the respective remodeling phases (purple = resorption, green = translation, orange = formation). (\**p*<0.05, \*\**p*<0.01, \*\*\**p*<0.001, \*\*\*\**p*<0.0001). Abbreviations: poly-aspartic acid (pAsp), non-mineralized (NM), mineralized (M), tartrate resistant acid phosphatase (TRAP), lactate dehydrogenase (LDH), micro-computed tomography (*μ*CT), day (D). The bone remodeling cycle illustration was modified from Servier Medical Art, licensed under a Creative Common Attribution 3.0 Generic License (http://smart.servier.com/, accessed on 8 July 2021).

As osteoclasts finished resorption around day 28, LDH activity as a measure for cell death, increased from day 21 to day 35 (Figure 6C). It is well accepted that differentiated osteoclasts have a relative short lifespan of about 2-3 weeks ^[42,43]^. Cell death after 21 days was therefore in line with our expectations. The increased osteoclastic activity in mineralized co-cultured constructs was however not associated with prolonged osteoclast survival (Figure 6C). On day 21 and 28, a higher LDH activity was even measured in cell supernatants of mineralized co-cultured constructs, indicating more cell death in these constructs. This was confirmed by DNA quantification at day 21 and 42, although not statistically significant (Figure 7Q). From day 21 on, osteogenic medium was provided which resulted in a further increase in mineralized volume in both scaffolds (Figure 6D+G-H). This increase was higher at all time points for mineralized scaffolds. In addition, formation sites in mineralized co-cultured constructs were localized over the entire scaffold and included spots which were previously resorbed, which might be attributed to osteoclast-osteoblast coupling (Figure 6H, yellow arrows and Figure S6). This could also explain the decrease in the cumulative percentage resorbed scaffold from day 21; resorption sites might have been filled with newly formed mineral (Figure 6E). Taken together, we were able to track the remodeling dynamics in our *in vitro* human bone model and these dynamics seemed to recapitulate the physiological bone remodeling cycle (Figure 6F).

**Figure 7.**
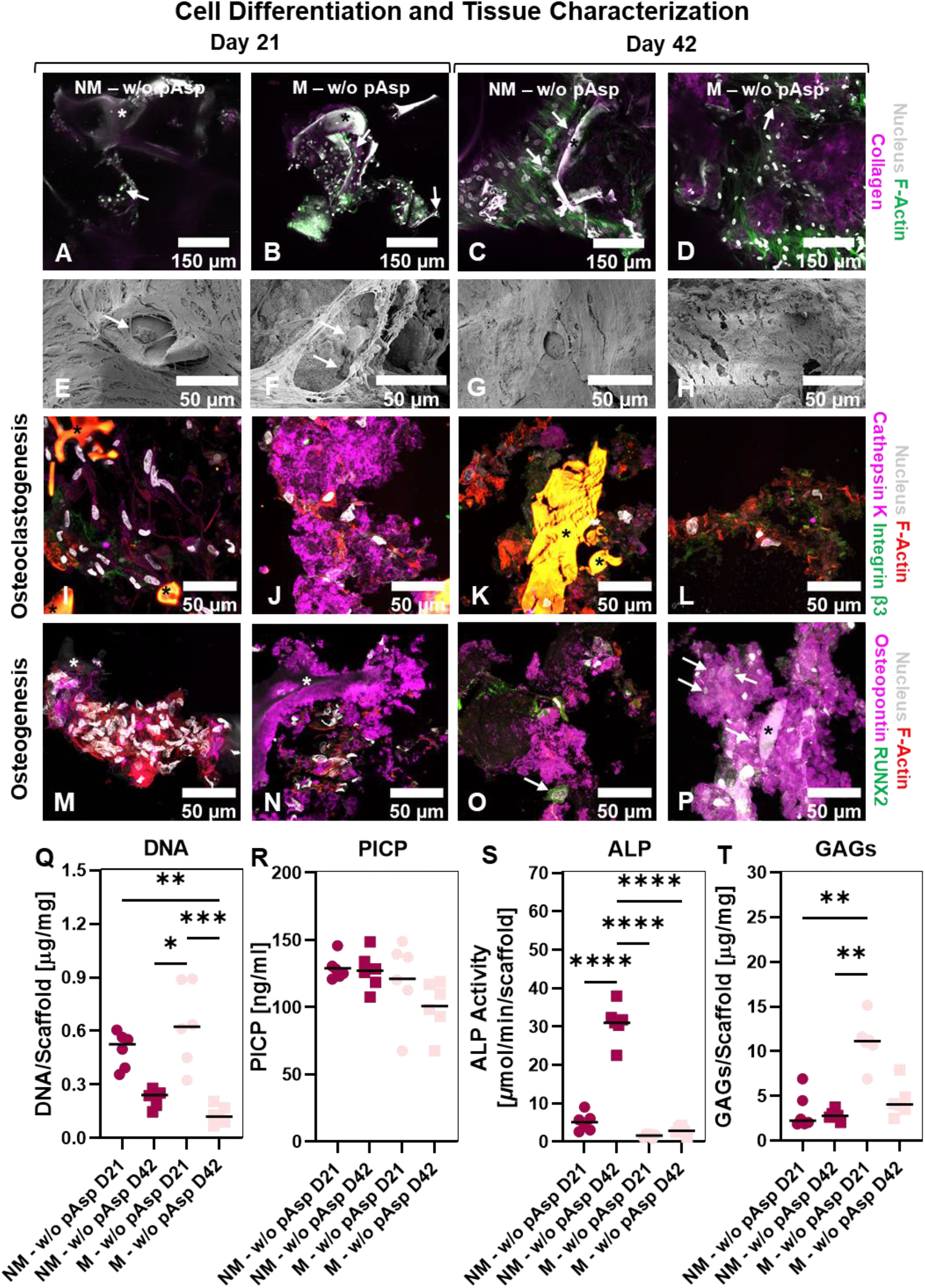
Cell differentiation and tissue formation in *in vitro* bone remodeling model. (**A-D**) Micrographs of 3D remodeling constructs, stained for collagen (magenta), F-Actin (green), and the nucleus (gray). White arrows indicate osteoclasts. (**E-H**) Morphology of and resorption sites on co-cultured constructs visualized with SEM. White arrows indicate osteoclasts. (**I-L**) Immunohistochemical analysis of sections for F-Actin (red), the nucleus (gray), and osteoclast markers cathepsin K (magenta) and integrin-*β*3 (green). (**M-P**) Immunohistochemical analysis of sections for F-Actin (red), the nucleus (gray), and osteogenic markers osteopontin (also produced by osteoclasts, magenta) and RUNX2 (green). Asterisks indicate the scaffold trabeculae. (**Q**) DNA quantification in co-cultured constructs, *p*<0.05 (Kruskal-Wallis and Dunn’s post hoc tests). (**R**) PICP quantification as a measure for collagen formation in co-cultured constructs, *ns* (One-way ANOVA). (**S**) ALP activity quantification as a measure for osteoblast activity, *p*<0.05 (One-way ANOVA and Holm-Šídák’s post hoc tests). (**T**) GAG content quantification, *p*<0.05 (Kruskal-Wallis and Dunn’s post hoc tests). (\**p*<0.05, \*\**p*<0.01, \*\*\**p*<0.001, \*\*\*\**p*<0.0001). Abbreviations: poly-aspartic acid (pAsp), non-mineralized (NM), mineralized (M), day (D), runt-related transcription factor 2 (RUNX2), pro-collagen 1 c-terminal propeptide (PICP), alkaline phosphatase (ALP), glycosaminoglycan (GAG).

Next, we characterized cell differentiation and organic matrix formation by the cells in the model. The presence of multinucleated cells was confirmed at day 21 and day 42 for both conditions by staining of the nucleus and the cytoskeleton (Figure 7A-D, white arrows). Osteoclast-like cells were also observed on the scaffold surface after 21 days of culture with SEM (Figure 7E+F, white arrows). These cells seem to resorb the mineral surface on mineralized scaffolds (Figure 7F). The osteoclast resorption marker cathepsin K was also highly expressed by cells on mineralized scaffolds in the resorption phase, indicating more functional osteoclasts on these scaffolds (day 21, Figure 7J). Interestingly, an excess of osteopontin, which can be produced by both osteoclasts and osteoblasts, was found in mineralized constructs after 21 days of culture. *In vivo*, osteopontin is also found on mineralized surfaces and is a major component of the cell-matrix interface (cement line) ^[44]^. Osteoclastic osteopontin is important for sealing zone formation and osteoclast migration ^[45–47]^. On mineralized scaffolds, osteoclasts have likely secreted osteopontin to allow for attachment and subsequent resorption ^[47]^. Osteopontin is also known as mineralization inhibitor in its phosphorylated state ^[48]^. However, excessive mineralization in mineralized scaffolds was still observed, meaning that the amount of osteopontin was not sufficient or that osteopontin was de-phosphorylated by osteoclasts through TRAP ^[48]^. After 42 days, osteopontin was present in both conditions and osteogenic differentiation was confirmed by the presence of nuclear runt-related transcription factor 2 (RUNX2). In addition, little collagen formation was observed at both time-points and in both conditions (Figure 7A-D), but mostly in the mineralized co-cultured constructs after 42 days of culture (Figure 7D). By measuring PICP in the medium, collagen formation at day 21 and 42 was quantified. Collagen type 1 formation was comparable for non-mineralized and mineralized co-cultured constructs (Figure 7R). While osteogenic differentiation medium was supplied from day 21, no further increase in collagen synthesis was observed. Collagen synthesis even tended to decrease in mineralized co-cultured constructs on day 42. This might be explained by a lack of mechanical loading in the system, which is crucial for *in vivo* bone adaptation and *in vitro* woven bone formation including collagen synthesis ^[49,50]^. Another explanation might be the excessive mineralization upon osteogenic stimulation in mineralized co-cultured constructs (Figure 6D). Mineralization occurred over the entire scaffold surface (Figure S6). As such, remodeling might have been terminated and cells therefore have undergone apoptosis, have been terminally differentiated into quiescent bone lining cells, or have been embedded into mineralized matrix and differentiated into osteocytes ^[51]^. This could also explain the differences found in ALP activity from the construct lysates (Figure 7S). Cells in non-mineralized constructs have clearly undergone differentiation towards ALP producing and thus mineralizing osteoblasts. As *in vitro* mineralization with osteogenic differentiation medium occurs after dephosphorylation of *β-* glycerophosphate by ALP, it is expected that the increase in mineralization for mineralized constructs was the result of ALP synthesis by the cells in these constructs. This could however not be detected on day 42, underlining the hypothesis that remodeling has been terminated on the mineralized SF scaffolds ^[52]^. Interestingly, a statistically significant higher sulphated GAG content was found on day 21 (Figure 7T). These GAGs were visualized between the trabecular-like structures (Figure S7). Although the origin of these GAGs is unclear, they have been shown to inhibit collagen degradation by osteoclastic cathepsin K, promote osteogenic differentiation and bone-like matrix formation, and promote mineralization ^[53–56]^.

## 3. Discussion

Current *in vitro* 3D bone remodeling models often lack the spatiotemporal investigation of the remodeling events *(i.e.* resorption, transition, formation) by starting their culture with osteoblast (progenitors) ^[11,12]^, or by only looking at osteoclast and osteoblast markers with *e.g.* gene expression or enzymatic activity assays rather than at their functionality to resorb and form a bone-like matrix ^[13,14]^. To enable the investigation of functional cell-matrix interactions and their spatiotemporal dynamics, materials should be developed that support osteoclast and osteoblast functionality. Therefore, we developed a bioinspired scaffold using SF as fibrous organic protein, that was mineralized with hydroxyapatite under influence of the non-collagenous protein mimic pAsp.

Like as in collagen, pAsp in the mineralization solution was instrumental for mineral infiltration into the SF. As a result, minerals appeared inside and on the surface of the SF films and trabecular-like structures within the scaffolds. This is comparable to bone, where minerals appear inside (intrafibrillar) and outside (extrafibrillar) to the collagen fibrils ^[20]^. In collagen, pAsp functions as a mineralization inhibitor in solution; guiding minerals to the collagen gap-region where the confinement induces mineral nucleation ^[57]^. Here, we have shown that pAsp functions similarly for the mineralization of SF, although the mineral nucleation mechanism is still unclear. One reasonable possibility is that the hydrophilic regions in SF allow for amorphous calcium phosphate infiltration and confinement, resulting in mineralization associated with the SF ^[24]^. When merged into the SF, pAsp could not improve mineral distribution through the scaffold. The addition of pAsp to the material even seemed to induce more heterogeneous mineralization (chunks instead of a layer). Twice earlier (to our knowledge), the influence of SF – pAsp materials on mineralization has been studied ^[58,59]^. Comparable to our results, Kim et al. (2008) ^[59]^ found chunks of mineral on the surface of their scaffold when ~ 9 wt% pAsp was added to the SF. Ma et al., (2010) ^[58]^ also found a comparable result for SF with 5% pAsp added. The minerals became more homogeneous when higher concentrations were added (i.e. 15%) ^[58]^. As there was no clear cytotoxic effect of the addition of pAsp to the SF in our study, the addition of higher concentrations of pAsp needs to be investigated. This might further increase the hydrophilicity and therefore also improve cell proliferation, osteoprogenitor differentiation, and osteoclastic resorption as reported in literature ^[34–37]^. These effects where not observed with the addition of 5% pAsp. However, the stiffness is expected to decrease further with higher concentrations of pAsp which could affect cell behavior ^[60]^. To further improve the mineral distribution through large scaffolds, perfusion of the mineralization solution needs to be explored ^[61]^.

Other *in vitro* remodeling models have used synthetic (mineralized) polymers, organic matrices or inorganic materials in the form of hydrogels or woven scaffolds ^[14]^. When composite materials are used for *in vitro* remodeling studies, organic materials are often mineralized by blending it with inorganic salts during fabrication or by coating it with supersaturated solutions ^[62,63]^. As such, most 3D materials used for current *in vitro* remodeling models lack mimicry with physiological bone. While biomineralized collagen type 1 scaffolds, featuring all components of physiological bone, are a promising material for *in vitro* bone remodeling models, they are often difficult to fabricate at the high density found in physiological bone. Recently, researchers looked at osteoclastic differentiation and resorption on such scaffolds. They found that despite its biomimicry, osteoclasts were unable to resorb the scaffold, probably as a result of the low fiber density ^[64]^. Here, we found that mineralized SF could support osteoclastic resorption. Mineralized SF scaffolds also seemed to stimulate a more physiologically relevant cell phenotype, indicated from their cathepsin K, osteopontin and glycosaminoglycan synthesis. In addition, using SF rather than collagen type 1 allows for a differential analysis of supplied and formed material ^[50]^. As altered collagen type 1 formation is a hallmark for bone pathologies like osteogenesis imperfecta and osteoporosis ^[65,66]^, the ability to study its formation should be considered for *in vitro* bone models ^[17]^. One limitation of the use of solely composite materials is likely the reduced osteoclast-osteoblast coupling. *In vivo*, coupling includes besides secreted, cell-bound, and topographical ques, also the release of growth factors from the bone matrix ^[67]^. Factors like transforming growth factor *β* (TGF-*β*), bone morphogenetic protein 2 (BMP-2), platelet derived growth factor (PDGF), and insulin-like growth factor (IGF) that are deposited by osteoblasts, stored in the matrix and released upon resorption, potentially stimulate MSC migration and osteogenic differentiation ^[67]^. While the use of native bone matrix could facilitate full investigation of cell-matrix and cell-cell interactions, proper decellularization needs to be performed as osteocyte (90% of bone cell population) apoptosis could induce pathological osteoclastic resorption ^[68]^. Introducing a pre-model phase where bone-like matrix is built by osteoblasts before remodeling is initiated might overcome these limitations ^[11,69]^. However, these models are time consuming, laborious, and might face reproducibility issues as the to be remodeled matrix is already susceptible for variation. Such complex models might improve the mimicry to bone remodeling *in vivo* but might in parallel complicate drug screening *in vitro*.

In the *in vitro* model presented in this study, coupling was observed by re-mineralization of resorption sites after osteogenic medium was provided. Mineralization was however not limited to resorption sites and the total mineralized volume was therefore increased over time. As healthy bone remodeling is characterized by balanced resorption and formation, our model does probably not yet fully represent the homeostatic physiological bone remodeling environment. While mineral resorption and formation was unbalanced in mineralized scaffolds, collagen synthesis as part of the osteoblastic formation seemed to stay behind with mineralization. Only little collagen formation could be detected in our model, despite the presence on osteogenic differentiation factors after 42 days. *In vivo*, the bone formation phase takes about 4 – 5 months and starts with osteoid *(i.e.* collagen and non-collagenous proteins) formation followed by mineralization ^[70]^. Most likely, the addition of exogenous phosphate with the *β*-glycerophosphate supplement in osteogenic medium steers this balance towards mineralization with limited osteoid formation and thus osteoblastic control ^[17]^. As in our model osteoclasts dissolved mineral from the scaffold, phosphate might have been released into the medium and the additional supplementation with *β*-glycerophosphate might have been redundant. The influence and the optimization of environmental factors (*e.g.* supplied medium or mechanical loading) should therefore be considered for future studies. For example, applying fluid shear stress to the cells to stimulate the osteoid formation and thereby potentially improving osteoblastic control over mineralization [^17,50,71,72]^.

## 4. Conclusion

Taken together, we have successfully exploited collagen mineralization techniques to mineralize SF films and scaffolds. In this regard, pAsp was instrumental to guide minerals into the SF structure. Mineralized SF scaffolds have subsequently demonstrated to support osteoclastic differentiation and resorption and to enhance mineralization. Functional cell-matrix interactions and their dynamics were successfully tracked with mainly non-destructive methods *(μCT* and medium analyses). The observed remodeling dynamics recapitulated the physiological bone remodeling cycle. Therefore, our *in vitro* bone remodeling model may reduce animal experiments and advance *in vitro* drug development for bone remodeling pathologies like osteoporosis where cell-matrix interactions need to be targeted.

## 5. Experimental Section

### 5.1 Preparation of silk fibroin films and scaffolds

Bombyx mori L. silkworm cocoons were degummed by boiling them in 0.2 M Na2CO3 for 1 h. After drying, silk was dissolved in 9 M LiBr, filtered, and dialyzed against UPW for 36 h using SnakeSkin Dialysis Tubing (11532541, Thermo Fisher Scientific, Breda, The Netherlands). After dialysis, the mass fraction of SF in solution was determined by measuring the dry weight per ml SF solution after lyophilization. For SF w/5% pAsp films and scaffolds, 5 wt% poly-aspartic acid sodium salt (P3418, Sigma-Aldrich, Zwijndrecht, The Netherlands) was mixed into the dialyzed SF solution. SF solution was then frozen at −80° C and lyophilized for 7 days. Lyophilized SF and SF with 5 wt% pAsp were dissolved in hexafluoro-2-propanol (003409, Fluorochem, Hadfield, UK) at a concentration of 17% (w/v) and casted onto 10 mm diameter cover slips (for SF films), or in scaffold molds containing NaCl granules with a size of 250-300 μm as template for the pores (for SF scaffolds). Hexafluoro-2-propanol in SF films was directly allowed to evaporate for 3 days. Scaffold molds were first covered to improve the SF blending with the granules. After 3 h, covers were removed and hexafluoro-2-propanol was allowed to evaporate for 7 days. After complete evaporation, *β*-sheets were induced by submerging SF films and SF-salt blocks in 90% MeOH for 30 min. NaCl was dissolved from the scaffolds in UPW, resulting in porous sponges. These sponges were cut into scaffolds of 3 mm in height and 5 mm in diameter.

### 5.2 Mineralization treatment

For mineralization of scaffolds and films, a 10x SBF stock was prepared as described by A.C. Tas and S.B. Bhaduri (2004) ^[27]^. Just prior to mineralization, mineralization solution was prepared by adding 100 μg/ml pAsp to 10x SBF, followed by the addition of NaHCO3 until a final concentration of 10 mM, both under vigorous steering. This resulted in a mineralization solution with a pH of ~6.3. For 10X SBF controls, pAsp was not added to the mineralization solution. Films and scaffolds were incubated for 2 weeks at 37 °C on an orbital shaker at 150 RPM in mineralization solution with a solution replenishment after 1 week. Mineralization solution volume was calculated from the apparent surface area of the sample as described by T. Kokubo and H. Takadama (2006) [^26^], where r is the radius of the sample and h the height (Equation 1). SF films were considered 2D.

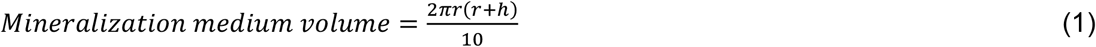

After mineralization, scaffolds and films were washed 3 x 15 min in an excess of UPW. Films and scaffolds for cell experiments were sterilized by autoclaving in phosphate buffered saline (PBS) at 121° C for 20 min.

### 5.3 Cell culture

#### 5.3.1 Monocyte isolation

Peripheral blood mononuclear cells (PBMCs) were isolated from a human peripheral blood buffy coat (Sanquin, Eindhoven, The Netherlands) of one healthy donor after written informed consent per the Declaration of Helsinki and according to the institutional guidelines. The buffy coat (~50 ml) was diluted with 0.6% w/v sodium citrate in PBS (citrate-PBS) until a final volume of 200 ml and layered per 25 ml on top of 10 ml Lymphoprep™ (07851, StemCell technologies, Köln, Germany) in 50 ml centrifugal tubes. After density gradient centrifugation (20 min at 800x g, lowest break), PBMCs were collected, resuspended in citrate-PBS, and washed four times in citrate-PBS supplemented with 0.01% bovine serum albumin (BSA, 10735086001, Sigma-Aldrich). PBMCs were frozen at 10^5^ cells/ml in freezing medium containing RPMI-1640 (RPMI, A10491, Thermo Fisher Scientific), 20% fetal bovine serum (FBS, BCBV7611, Sigma-Aldrich) and 10% dimethyl sulfoxide (DMSO, 1.02952.1000, VWR, Radnor, PA, USA) and stored in liquid nitrogen until further use. Before MC isolation, PBMCs were thawed, collected in medium containing RPMI, 10% FBS (BCBV7611, Sigma-Aldrich) and 1% penicillin-streptomycin (p/s, 15070063, Thermo Fisher Scientific), and after centrifugation resuspended in isolation buffer (0.5% w/v BSA in 2mM EDTA-PBS). MCs were enriched from PBMCs with manual magnetic activated cell separation (MACS) using the Pan Monocyte Isolation Kit (130-096-537, Miltenyi Biotec, Leiden, Netherlands) and LS columns (130-042-401, Miltenyi Biotec) according to the manufacturer’s protocol, and directly used for experiments.

#### 5.3.2 Mesenchymal stromal cell isolation and expansion

MSCs were isolated from human bone marrow (Lonza, Walkersville, MD, USA) and characterized for surface markers and multilineage differentiation, as previously described ^[73]^. MSCs were frozen at passage 4 with 1.25*10^6^ cells/ml in freezing medium containing FBS (BCBV7611, Sigma-Aldrich) with 10% DMSO and stored in liquid nitrogen until further use. Before experiments, MSCs were thawed, collected in high glucose DMEM (hg-DMEM, 41966, Thermo Fisher Scientific), seeded at a density of 2.5*10^3^ cells/cm^2^ and expanded in medium containing hg-DMEM, 10% FBS (BCBV7611, Sigma-Aldrich), 1% Antibiotic Antimycotic (anti-anti, 15240, Thermo Fisher Scientific), 1% Non-Essential Amino Acids (11140, Thermo Fisher Scientific), and 1 ng/mL basic fibroblastic growth factor (bFGF, 100-18B, PeproTech, London, UK) at 37 °C and 5% CO_2_. After 9 days, cells were detached using 0.25% trypsin-EDTA (25200, Thermo Fisher Scientific) and directly used for experiments at passage 5.

#### 5.3.3 Two-dimensional monocyte and mesenchymal stromal cell mono-cultures

For 2D MC and MSC mono-cultures, films were pre-wettened overnight at 37 °C in osteoclast control medium (α-MEM (41061, Thermo Fisher Scientific), 10% human platelet lysate (hPL, PE20612, PL BioScience, Aachen, Germany) and 1% anti-anti) and osteogenic control medium (lg-DMEM (22320, Thermo Fisher Scientific), 10% hPL and 1% anti-anti). Before seeding, medium was removed, and cells were seeded by pipetting 5 μl of cell suspension (1.5*10^5^ cells/5 μl for MCs and 2.5*10^4^ cells/5 μl for MSCs) onto the films. Cells were allowed to attach for 90 min at 37 °C and every 20 minutes a small droplet of the respective control medium was added to prevent for drying of the films. MCs were first cultured in priming medium (osteoclast control medium + 50 ng/ml macrophage colony-stimulating factor (M-CSF, 300-25, PeproTech)). After 48 hours, priming medium was replaced by osteoclast medium (priming medium + 50 ng/ml receptor activator of nuclear factor κB ligand (RANKL, 310-01, PeproTech)) to induce osteoclastic differentiation. MSCs were stimulated to undergo osteogenic differentiation with osteogenic medium (osteogenic control medium + 10 mM *β-* glycerophosphate (G9422, Sigma-Aldrich), 50 μg/ml ascorbic acid-2-phosphate (A8960, Sigma Aldrich), and 100 nM dexamethasone (D4902, Sigma-Aldrich)). Cells were kept in culture for 7 days at 37 °C and 5% CO_2_, medium was replaced on day 2 and 5 and medium samples were collected and stored at −80 °C. Films were sacrificed for analyses after 2 days and 7 days of culture.

#### 5.3.4 Three-dimensional monocyte-mesenchymal stromal cell co-culture

Scaffolds were pre-wettened overnight at 37 °C in osteoclast control medium. Before seeding, medium was removed, and cells were resuspended in osteoclast control medium (2.5*10^6^ MCs and 5*10^5^ MSCs/20 μl) and seeded by pipetting 20 μl of cell suspension onto the scaffolds. Cells were allowed to attach for 90 min at 37 °C and every 20 minutes a small droplet of osteoclast control medium was added to prevent for drying of the scaffolds. The cell-loaded scaffolds were statically cultured for 6 weeks at 37 °C and 5% CO_2_ in custom-made bioreactors, which allowed for *μ*CT scanning during the culture period. Cells were cultured in osteoclast medium for the first 3 weeks (priming medium for the first 48 h whereafter medium was replaced by osteoclast medium). After 3 weeks, medium was switched to osteogenic medium to stimulate osteogenic differentiation. Medium was replaced 3x per week and medium samples were collected weekly and stored at −80 °C. Constructs were sacrificed for analyses after 3 weeks (day 21) and after 6 weeks (day 42) of culture.

### 5.4 Analyses

#### 5.4.1 Contact angle measurements

Water contact angles were measured for SF films w/o pAsp and w/5% pAsp on a Dataphysics OCA30 contact angle goniometer *(N* = 5 per group). A 2 μl droplet of UPW was deposited on the films and after approximately 2 s the contact angles were determined by fitting the contour of the droplet using OCA20 software.

#### 5.4.2 Mineral precipitation in medium

Mineralization solution samples were collected from mineralized films after 1 week and 2 weeks of mineralization (*N* = 8 per condition). Mineral precipitation in the mineralization solution was determined by measuring the optical density of 100 μl sample in a 96-wells assay plate at 600 nm using a plate reader (Synergy™ HTX, Biotek).

#### 5.4.3 Calcium assay

Films (*N* = 5 per condition) were lyophilized and incubated for 48 h in 5 wt% trichloroacetic acid (TCA, T6399, Sigma-Aldrich). Scaffolds (*N* = 5 per condition) were lyophilized, weighted, disintegrated in 5 wt% trichloroacetic using 2 steel balls and a mini-beadbeater^TM^ (Biospec, Bartlesville, OK, USA), and subsequently incubated for 48 h. After incubation, a calcium assay (Stanbio, 0150-250, Block Scientific, Bellport, NY, USA) was performed to quantify calcium content in both films and scaffolds according to the manufacturer’s instructions. Briefly, 95 μl Cresolphthalein complexone reaction mixture was added to 5 μl sample and incubated at room temperature for 1 min. Absorbance was measured at 550 nm with a plate reader and absorbance values were converted to calcium concentrations using standard curve absorbance values.

#### 5.4.4 Mechanical analyses

Mechanical tests of films (*N* = 5 per condition) were performed with a Piuma nanoindenter (Optics 11, Amsterdam, The Netherlands) equipped with a spherical indenter tip probe with a radius of 29.1 μm and a stiffness of 204.6 N/m (p190853, Optics 11). Films were tested in PBS and an indentation of 10 μm depth was performed at 4 random locations per film and the Young’s modulus was derived by fitting the load-depth curves to the Hertzian contact model between 0% and 30% of the maximum load point, assuming a Poisson’s ratio of 0.4 ^[74,75]^. Scaffolds (*N* = 5 per condition) were mechanically tested in PBS by a full unconfined compression test using a 500 N load cell on a Criterion 42 mechanical test system (MTS, Berlin, Germany). Samples were compressed at a rate of 17% displacement/min until a displacement of 60% from the sample height was reached. The Young’s modulus was derived by a linear fit to the load-displacement curves between 2% and 10% displacement using MATLAB (version 2019b, The MathWorks Inc., Natrick, MA, USA).

#### 5.4.5 Scanning electron microscopy (SEM)

Samples *(N* = 3-4 per experiment, time point, and condition) were fixed in 2.5% glutaraldehyde in 0.1 M sodium cacodylate buffer (CB) for 4 h and then washed in CB. For the characterization of (mineralized) scaffolds, both 3D samples and cross-sections were prepared. For cross-sections, scaffolds were after fixation soaked for 15 minutes in each 5% (w/v) sucrose and 35% (w/v) sucrose in PBS. Scaffolds were embedded in Tissue Tek® (Sakura) and frozen with liquid N_2_. Cryosections were prepared with a thickness of 5 μm on 10 x 10 mm indium tin oxide (ITO) coated glass slides (576352, Sigma-Aldrich). Tissue Tek® was removed by washing with distilled water. Co-cultured scaffolds (*N* = 2 out of 4 per time point and condition) were stained and imaged with confocal microscopy as described below before dehydration. All samples were dehydrated with graded ethanol series (37%, 67%, 96%, 3 x 100%, 15 minutes each), followed by a hexamethyldisilazane (HDMS)/ethanol series (1:2, 1:1, 2:1, 3 x 100% HDMS, 15 minutes each). Samples were coated with 20 nm gold and imaging was performed in high vacuum, at 10 mm working distance, with a 5kV electron beam (Quanta 600F, FEI, Eindhoven, The Netherlands).

#### 5.4.6 Alizarin red

Alizarin red staining was performed on films, cross-sections of films, and cross-sections of scaffolds (*N* = 3 per experiment and condition). Samples were fixed overnight in 3.7% neutral buffered formaldehyde and washed twice with PBS. Samples for cross-sections were prepared as described above (Section SEM) and cryosections were sliced with a thickness of 5 μm on Epredia™ SuperFrost Plus™ Adhesion slides (Fisher Scientific, Breda, The Netherlands). Samples were washed in distilled water and stained for 15 min in 2% w/v Alizarin Red (ab146374, Abcam, Cambridge, UK) in distilled water at a pH of ~4.2. Films were directly washed in distilled water and imaged upon staining. Cross-sections were first dehydrated in pure acetone, acetone/xylene (1:1) and pure xylene, and mounted with Entellan™ (1.07960, Sigma-Aldrich). Samples were imaged with bright field microscopy (Zeiss Axio Observer Z1 with a 20x/0.8 Plan-Apochromat objective or a 5x/0.13 EC Epiplan-Neofluar objective).

#### 5.4.7 Micro-computed tomography

For *μ*CT scanning, wet and dry mineralized scaffolds (*N* = 5 per condition) and co-cultured constructs (*N* = 8 per condition) were scanned and analyzed with a *μ*CT100 imaging system (Scanco Medical, Brüttisellen, Switzerland). Scanning was performed with an energy level of 45 kVp, intensity of 200 μA, integration time of 300 ms and with twofold frame averaging. To reduce part of the noise, a constrained Gaussian filter was applied to all scans with filter support 1 and filter width sigma 0.8 voxel. For mineralized scaffolds (both wet and dry), scanning was performed at an isotropic resolution of 11.4 μm. Filtered images were segmented, for wet scaffolds to detect mineralization (global threshold of 27% of the maximum grayscale value) and for dry scaffolds to study the morphology (global threshold of 22% of the maximum grayscale value). Unconnected objects smaller than 30 voxels were removed through component labeling. Morphology parameters were computed from dry scaffolds using the scanner manufacturer’s image processing language (IPL) ^[76]^. To determine the pore size distribution, the image background was filled with largest possible spheres of which the diameter was derived. To quantify the degree of connectivity between trabecular-like structures, the mean connectivity density was calculated per scaffold according to a previously described method ^[77]^. In addition, porosity, mean trabecular thickness, mean trabecular space and average trabecular number per mm were derived per scaffold after triangulation of segmented scaffolds using the plate model. To track mineralization, co-cultured scaffolds were scanned weekly after an initial baseline scan (day 2) at an isotropic resolution of 17.2 μm. Filtered scans were segmented at a global threshold of 24% of the maximum grayscale value and unconnected objects smaller than 30 voxels were removed through component labelling. In addition, follow-up images of the radiopaque mineralized co-cultured scaffolds were registered to baseline images such that voxels at the surface of the scaffold were categorized into resorption site, formation site, or unchanged site ^[78]^. The scaffold was segmented at a global threshold of 24% of the maximum grayscale value and remodeled scaffold surface was segmented at a global threshold of 7.5% of the maximum grayscale value, which was chosen after registration of cell-free construct images in such a way that resorption and formation were below ~1.5% of the total volume. To reduce noise, only a minimum cluster of 2 resorbed or formed voxels were included in the analyses, meaning that only resorption and formation sites of more than ~30 μm in length could be detected.

#### 5.4.8 Raman microscopy

Scaffold cross-sections were analyzed with Raman Microscopy. Scaffolds *(N* = 3) were soaked for 15 minutes in each 5% (w/v) sucrose and 35% (w/v) sucrose in phosphate buffered saline (PBS). Samples were embedded in Tissue Tek® (Sakura) and quickly frozen with liquid N_2_. Cryosections were prepared with a thickness of 10 μm on microscope glasses covered with aluminum foil. Sections were washed three times with distilled water and air dried. Raman microscopy was subsequently performed on a Witec Alpha 300 R instrument (Witec, Ulm, Germany). Spectra were obtained using a 457 nm excitation laser at 8 mW. The light was split through a 600 mm^-1^ grating resulting in a spectral resolution of 2.8 cm^-1^. Spectral imaging was performed at a resolution of 1 μm at an exposure time of 1 s. The obtained data were analyzed using the Witec Project 5 software (Witec). Samples were background corrected with the automatic shape function in the software, using shape size 400. Component analysis was subsequently performed, and the two or three major components were presented. The spectra are formed by averaging all the pixels containing the unique chemical signature. After extraction the data was transferred to Origin (Origin Pro 2021, OriginLab Corporation, Northhampton, MA, USA) where the spectra were normalized to the Amide I 1660 cm^-1^ peak for visualization.

#### 5.4.9 X-ray photoelectron spectroscopy measurements

XPS spectra were obtained of air dried scaffolds using a Thermo Scientific K-Alpha spectrometer (Thermo Fisher Scientific) equipped with a 180° double-focusing hemispherical analyzer with a 128-channel detector that uses an aluminum anode (Al Kα, 1486.7 eV, 72 W) and monochromatic, small-spot X-ray source. The survey scans used a pass energy of 200 eV and the atomic region scans 50 eV. The atom compositions were quantified from the survey spectra and the ratio of different carbon bonds were determined from the carbon region spectra using CasaXPS software (version 2.3.23).

#### 5.4.10 Biochemical content analyses

Lyophilized mono-cultured films (*N* = 5 per condition) and co-cultured constructs (*N* = 6 per time point and per condition) were digested overnight in papain digestion buffer (containing 100 mmol phosphate buffer, 5 mmol L-cystein, 5 mmol EDTA and 140 μg/ml papain (P4762, Sigma-Aldrich)) at 60 °C. DNA was quantified using the Qubit Quantification Platform (Invitrogen) with the high sensitivity assay, according to the manufacturer’s instructions. GAG content in co-cultured constructs was measured using a dimethyl methylene blue (DMMB) assay ^[79]^ with shark cartilage chondroitin sulfate (C4284, Sigma-Aldrich) as a reference. Absorbance was read at 540 nm and 595 nm using a plate reader. Absorbance values were subtracted from each other (540-595) and converted to GAG content using standard curve absorbance values.

#### 5.4.11 Lactate dehydrogenase activity

LDH activity was measured over time in cell supernatants of mono-cultured films (*N* = 5) and co-cultured constructs (*N* = 6 – 12 per condition, 3 samples per bioreactor containing 4 scaffolds). A 100 μl supernatant sample or NADH (10107735001, Sigma-Aldrich) standard was incubated with 100 μl LDH reaction mixture (11644793001, Sigma-Aldrich) in 96-wells assay plates. Absorbance was measured after 5, 10 and 20 min at 490 nm, and LDH activity was calculated between 5 and 20 min reaction, using standard curve absorbance values.

#### 5.4.12 PrestoBlue™ assay

Mono-cultured films were incubated with a 10% v/v PrestoBlue^TM^ (A13262, Thermo Fisher Scientific) in osteogenic (for MSCs) or osteoclast (for MCs) control medium solution for 1 h at 37 °C in the dark. Fluorescence was measured with a plate reader (excitation: 530/25 nm, emission 590/35 nm). Measured fluorescence was corrected for blank medium samples.

#### 5.4.13 Tartrate resistant acid phosphatase activity

TRAP was measured over time in cell supernatants of co-cultured constructs (*N* = 6 – 12 per condition, 3 samples per bioreactor containing 4 scaffolds). A 10 μl supernatant sample or p-nitrophenol standard was incubated with 90 μl p-nitrophenyl phosphate buffer (1 mg/ml p-nitrophenyl phosphate disodium hexahydrate (71768, Sigma-Aldrich), 0.1 M sodium acetate, 0.1% Triton X-100 and 30 μl/ml tartrate solution (3873, Sigma-Aldrich) in PBS) in 96-wells assay plates for 90 min at 37 °C. To stop the reaction, 100 μl 0.3 M NaOH was added. Absorbance was read at 405 nm using a plate reader and absorbance values were converted to TRAP activity (converted p-nitrophenyl phosphate in nmol/ml/min) using standard curve absorbance values.

#### 5.4.14 Alkaline phosphatase activity

Co-cultured constructs (*N* = 6 per time point and per condition) were washed in PBS and disintegrated using 2 steel balls and a mini-beadbeater^TM^ (Biospec, Bartlesville, OK, USA) in cell lysis buffer containing 0.2% (v/v) Triton X-100 and 5 mM MgCl_2_. ALP activity in cell lysates was determined by adding 20 μl of 0.75 M 2-amino-2-methyl-1-propanol (A65182, Sigma-Aldrich) to 80 μl sample in 96-wells assay plates. Subsequently, 100 μl substrate solution (10 mM p-nitrophenyl-phosphate (71768, Sigma-Aldrich) in 0.75 M 2-amino-2-methyl-1-propanol) was added and wells were incubated at room temperature for 15 minutes. To stop the reaction, 100 μl 0.2 M NaOH was added. Absorbance was measured with a plate reader at 450 nm and these values were converted to ALP activity (converted p-nitrophenyl phosphate in μmol/ml/min) using standard curve absorbance values.

#### 5.4.15 Pro-Collagen 1 C-Terminal Propeptide quantification

PICP as collagen formation product was quantified in cell supernatants of co-cultured constructs from day 21 and day 42 using an enzyme-linked immunosorbent assay (ELISA, MBS2502579, MyBioSource, San Diego, CA, USA) according to the manufacturer’s protocol. Samples were added to anti-human PICP coated microwells. After 90 min incubation at 37 °C, samples were replaced by biotinylated antibody solution followed by 60 min incubation at 37 °C. After thorough washing, HRP-conjugate solution was added, and plates were incubated for 30 min at 37 °C. Wells were again washed, and substrate reagent was added followed by 15 min incubation in the dark at 37 °C. To stop the reaction, stop solution was added and absorbance was measured at 450 nm in a plate reader. Absorbance values were converted to PICP concentrations using standard curve absorbance values.

#### 5.4.16 (Immuno)histochemical analyses

Mono-cultured films after 7 days of culture (*N* = 3 per condition) were stained with DAPI and Phalloidin to visualize cell nuclei and the actin cytoskeleton, respectively. In short, films were fixed in 3.7% neutral buffered formaldehyde for 15 min, permeabilized in 0.5% Triton X-100 in PBS for 10 min and blocked in 2% BSA in PBS for 30 min. Cells were incubated with 0.1 μg/ml DAPI (D9542, Sigma-Aldrich) and 50 pmol Atto 647-conjugated Phalloidin (65906, Sigma-Aldrich) in PBS for 1 h. As some films had a curved surface, z-stacks were taken with a confocal laser scanning microscope (Leica TCS SP8X, 20x/0.4 HC PL Fluotar L objective). After background removal, to reduce autofluorescence from SF, z-stacks were converted to maximum intensity projections using FiJi ^[80]^.

Co-cultured scaffolds (*N* = 2 per time point and per condition) that were fixed for SEM analysis, were washed in PBS, permeabilized for 30 min in 0.5% Triton X-100 in PBS and stained overnight with 1 μmol/mL CNA35-mCherry ^[81]^ at 4 °C to visualize collagen. After washing with PBS, samples were incubated for 1 h with 0.1 μg/ml DAPI and 50 pmol Atto 488-conjugated Phalloidin (49409, Sigma-Aldrich). Samples were washed and imaged in PBS and z-stacks were acquired with a confocal laser scanning microscope (Leica TCS SP8X, 20x/0.75 HC PL APO CS2 objective). Z-stacks were converted to maximum intensity projections using FiJi ^[80]^.

Co-cultured scaffolds (*N* = 4 per time point and per condition) were prepared for cryosections by soaking them for 15 minutes in each 5% (w/v) sucrose and 35% (w/v) sucrose in phosphate buffered saline (PBS). Samples were embedded in Tissue Tek® (Sakura) and quickly frozen with liquid N_2_. Cryosections were prepared with a thickness of 30 μm for antibody stainings and with a thickness of 5 μm for alcian blue staining. Upon staining, sections were fixed for 15 minutes in 3.7% neutral buffered formaldehyde and washed twice with PBS.

To visualize proteoglycan deposition, sections were stained in 1% w/v alcian blue (A5268, Sigma-Aldrich) in 3% acetic acid solution (pH 2.5) for 30 min. After washing in running distilled water for 5 min, sections were placed in Mayer’s Hematoxylin solution for 10 min and washed in tunning tap water for 10 min. All sections were dehydrated in one change of 70% and 96% EtOH, three changes of 100% EtOH, and two changes of xylene. Sections were mounted with Entellan (107961 Sigma-Aldrich) and imaged with a bright field microscope (Zeiss Axio Observer Z1, Plan-Apochromat 100x/1.40 objective).

To study osteogenic differentiation, sections were stained with DAPI, Atto 488-conjugated Phalloidin, RUNX2 and osteopontin. To study osteclastic differentiation, sections were stained with DAPI, Atto 647-conjugated Phalloidin, Cathepsin K, and integrin-*β*_3_. Briefly, sections were permeabilized in 0.5% Triton X-100 in PBS for 10 min and blocked in 10% normal goat serum in PBS for 30 min. Primary antibodies were incubated overnight at 4 °C om 1% normal goat serum in PBS, secondary antibodies were incubated with 1 μg/ml DAPI and 50 pmol Phalloidin in PBS for 1 h at room temperature. Antibodies are listed in Table S1. Z-stacks were acquired with a laser scanning microscope (Leica TCS SP8X, 63x/1.4 HC PL Apo CS2 objective). Z-stacks were converted to maximum intensity projections using FiJi ^[80]^.

### 5.5 Statistical analyses

Statistical analyses were performed, and graphs were prepared in GraphPad Prism (version 9.3.0, GraphPad, La Jolla, CA, USA) and R (version 4.1.2) ^[82]^. Data were tested for normality in distributions and equal variances using Shapiro-Wilk tests and Levene’s tests, respectively. When these assumptions were met, mean ± standard deviation are presented, and to test for differences, an independent t-test (for the comparison of two groups), one-way ANOVA followed by Holm-Šídák’s post hoc method with adjusted *p*-values for multiple comparisons (for the comparison >2 groups), or a two-way ANOVA followed by Turkey’s post hoc tests with adjusted p-value for multiple comparisons (for comparisons between groups over a period of time) were performed. Other data are presented as median ± interquartile range and were tested for differences with non-parametric Mann–Whitney U tests or Kruskal-Wallis tests with Dunn’s post hoc tests with adjusted p-value for multiple comparisons. With a p-value of <0.05 differences were considered statistically significant.

## 7. Acknowledgement

The authors thank Dewy van der Valk and Jasper Aarts for their help with the chemical characterization of the samples. The authors gratefully acknowledge the financial support for B.d.W. and S.H. by the research program TTW with project number TTW 016.Vidi.188.021 to S.H., which is financed by the Netherlands Organization for Scientific Research (NWO). R.v.d.M, N.S. and A.A. were supported by the European Research Council (ERC) Advanced Investigator grant (H2020-ERC-2017-ADV-788982-COLMIN). A.A. was supported by the NWO (VI.Veni.192.094).

## 8. Data availability statement

The data that support the findings of this study are available from the corresponding author upon reasonable request.

