## Supplementary Material for "Bioinspired Silk Fibroin Mineralization for Advanced *In Vitro* Bone Remodeling Models"

To check for the presence of poly aspartic acid (pAsp) in silk fibroin (SF) w/5% pAsp films, films were stained with the cationic dye alcian blue to allow for visualization of the negatively charged pAsp (Figure S1).

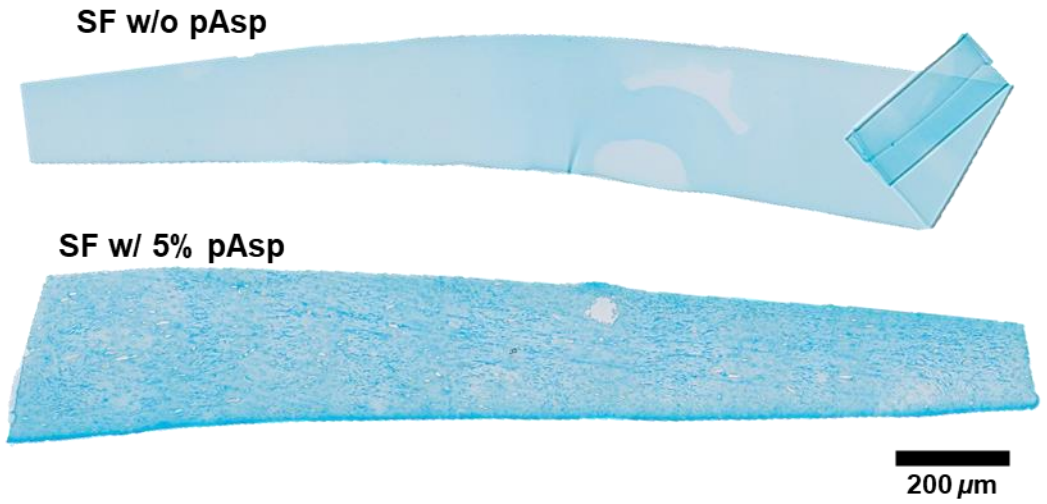


**Figure S1.** Plain silk fibroin (SF) film and SF film with 5 wt% poly-aspartic acid (pAsp) stained with alcian blue to visualize the negatively charged pAsp.

The presence of a small amount of pAsp in the SF material was also confirmed by chemical analysis. Raman spectroscopy measurements revealed a small peak at 1783 $cm^{-1}$, suggesting the presence of pAsp (Figure S2).

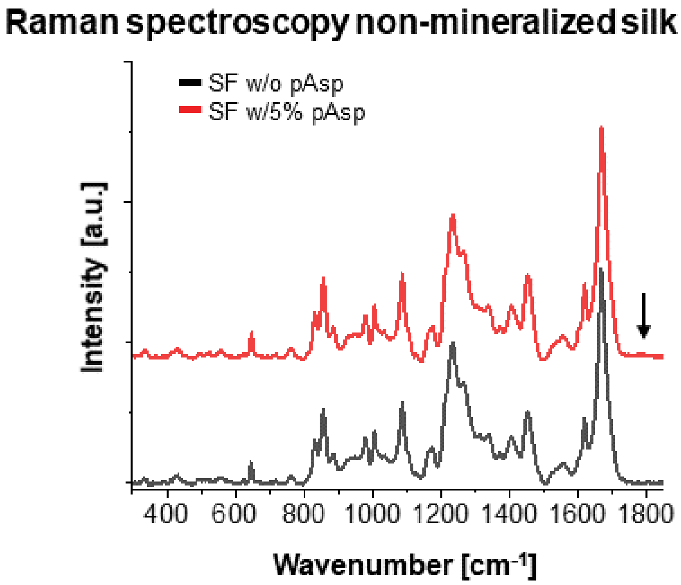


**Figure S2.** Comparison of non-mineralized silk fibroin (SF) in the presence (red) and absence (black) of poly-aspartic acid (pAsp). The spectra are near identical apart from a small peak at 1783 $cm^{-1}$, which indicates the presence of pAsp. Shown spectra are average spectra of a 30x30 µm area scans (total 900 spectra) for both samples.

X-ray photoelectron spectroscopy (XPS) measurements revealed a carbon peak with a wider shape, which is likely attributed to the carboxyl group in pAsp. XPS measurements also revealed the presence of calcium, phosphate and pAsp in both mineralized SF w/o pAsp and SF w/5% pAsp scaffolds. The presence of pAsp was observed by the carbon peak with wider shape relative to non-mineralized SF w/o pAsp scaffolds indicative for the presence of the carboxyl group of pAsp (Figure S3).

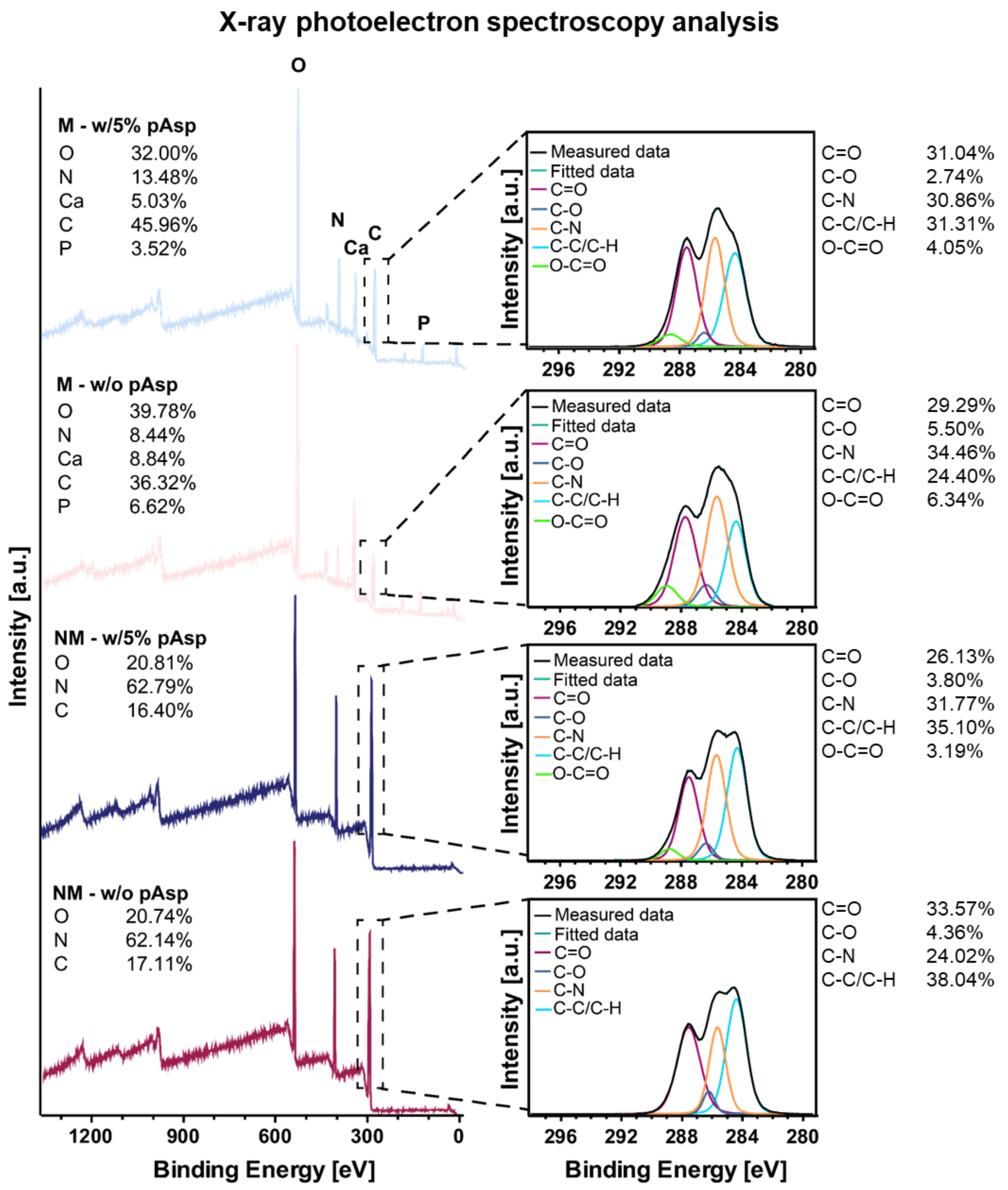

**Figure S3.** Comparison of X-ray photoelectron spectroscopy (XPS) measurements from mineralized and non-mineralized silk fibroin (SF) scaffolds w/o poly-aspartic acid (pAsp) and w/5% pAsp. Left panel presents the survey spectra with identified elements for each scaffold. Right panel present the carbon spectra, which were decomposed into the chemical bonds present in the different scaffolds.

To check for the presence of calcium in films and their cross-sections, an alizarin red staining was performed (Figure S4).

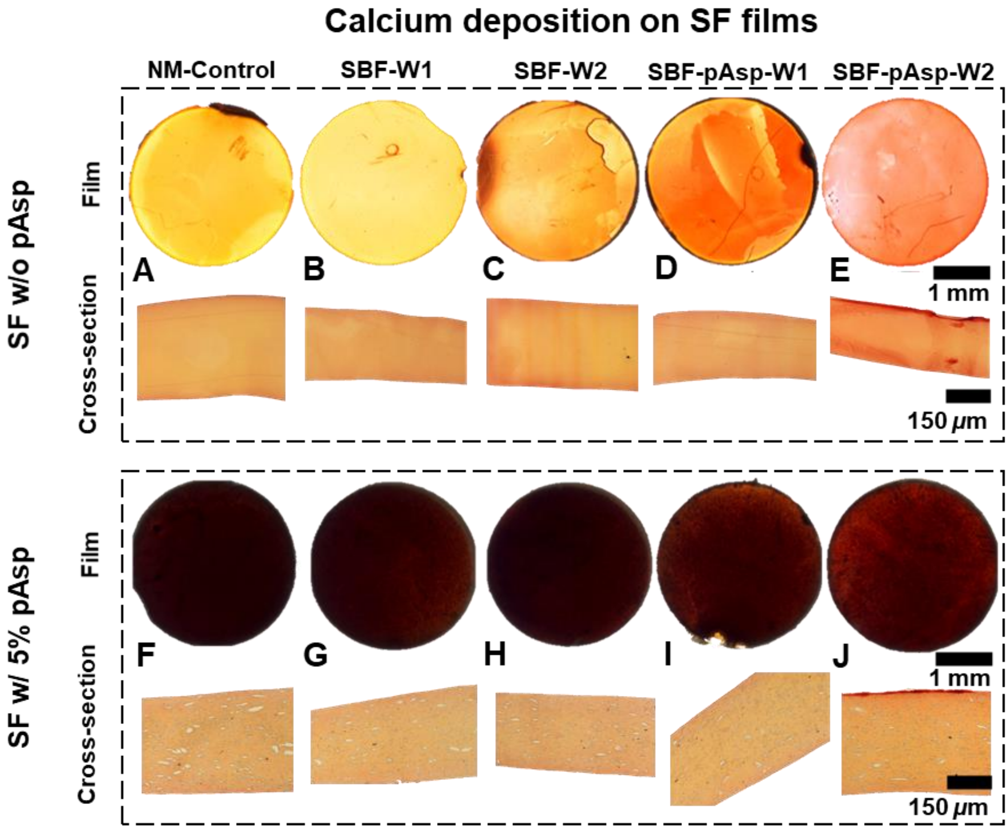


**Figure S4.** Calcium visualization with alizarin red staining. A clear red staining was observed on top of films mineralized with poly-aspartic acid (pAsp) in the mineralization solution after 2 weeks (cross-sections, **E+J**). In these groups, only mineralized silk fibroin (SF) w/o pAsp films showed red staining inside the film indicating mineral infiltration into the films (Figure S2E, cross-section). The films with pAsp in the material where less transparent and also non-mineralized (NM) films appeared to bind the stain. By preparing cross-sections, differences within this group became clearer. Abbreviations: week (W), simulated body fluid (SBF).

Because of the radiolucent nature of SF when immersed in water, mineralization could be localized with micro-computed tomography (µCT) scanning of the scaffolds. By drying the mineralized scaffolds, their 3D morphology could also be characterized after *µ*CT scanning (Figure S5).

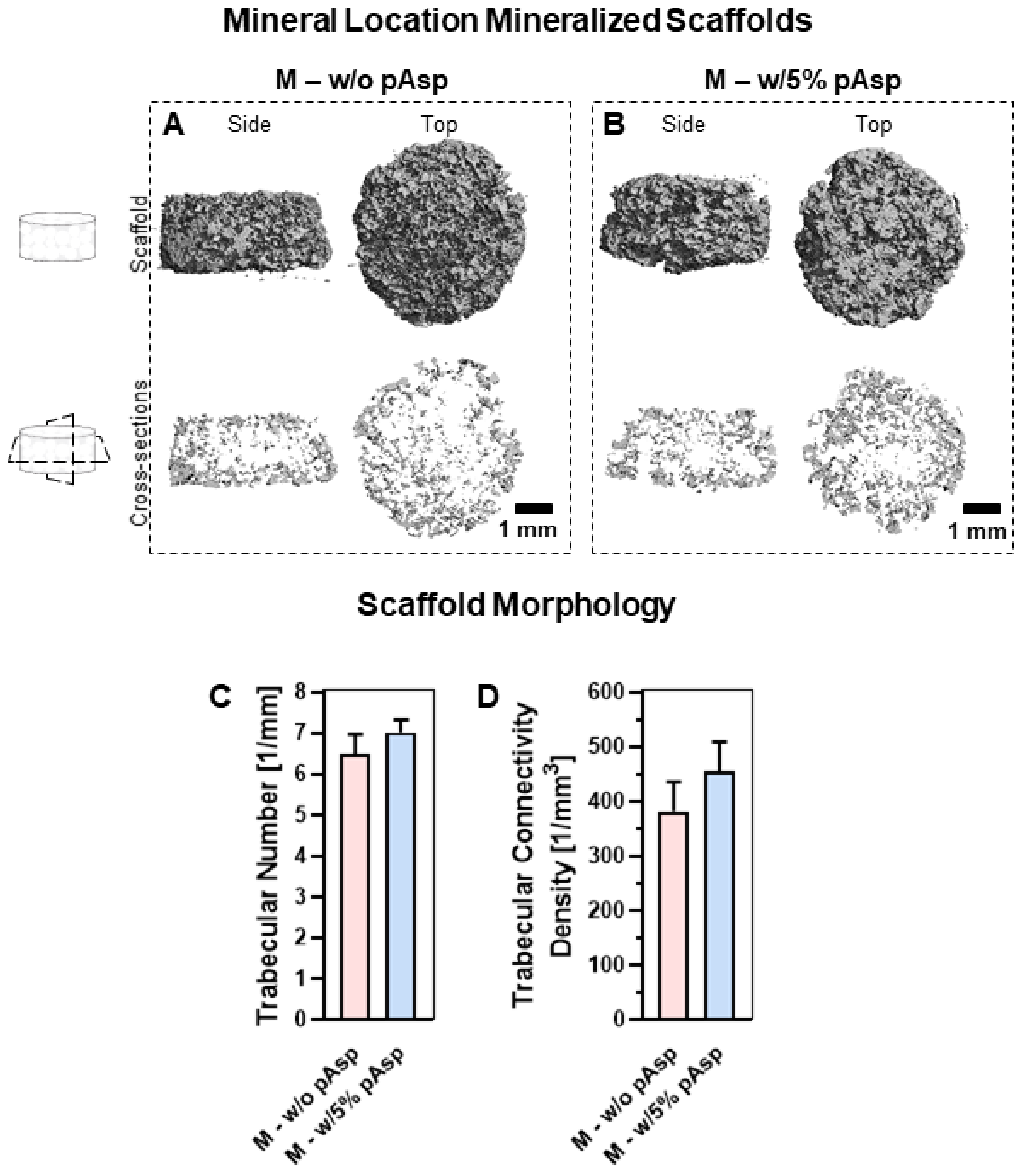


**Figure S5.** Morphological analyses of mineralized (M) silk fibroin (SF) scaffolds. (**A**) Mineral location visualized with *µ*CT for plain SF scaffolds and (**B**) for SF with 5% poly-aspartic acid (pAsp) in the scaffold. Cross-sections are optical slices of the whole scaffolds. No clear differences were observed between the two materials in terms of mineral distribution, (**C**) trabecular number (per mm length), and (**D**) the trabecular connectivity density, both *ns* (Independent t-test).

Longitudinal $\mu$CT scanning was performed to track the remodeling dynamics. The influence of scanning on cytotoxicity was also evaluated (Figure S6).

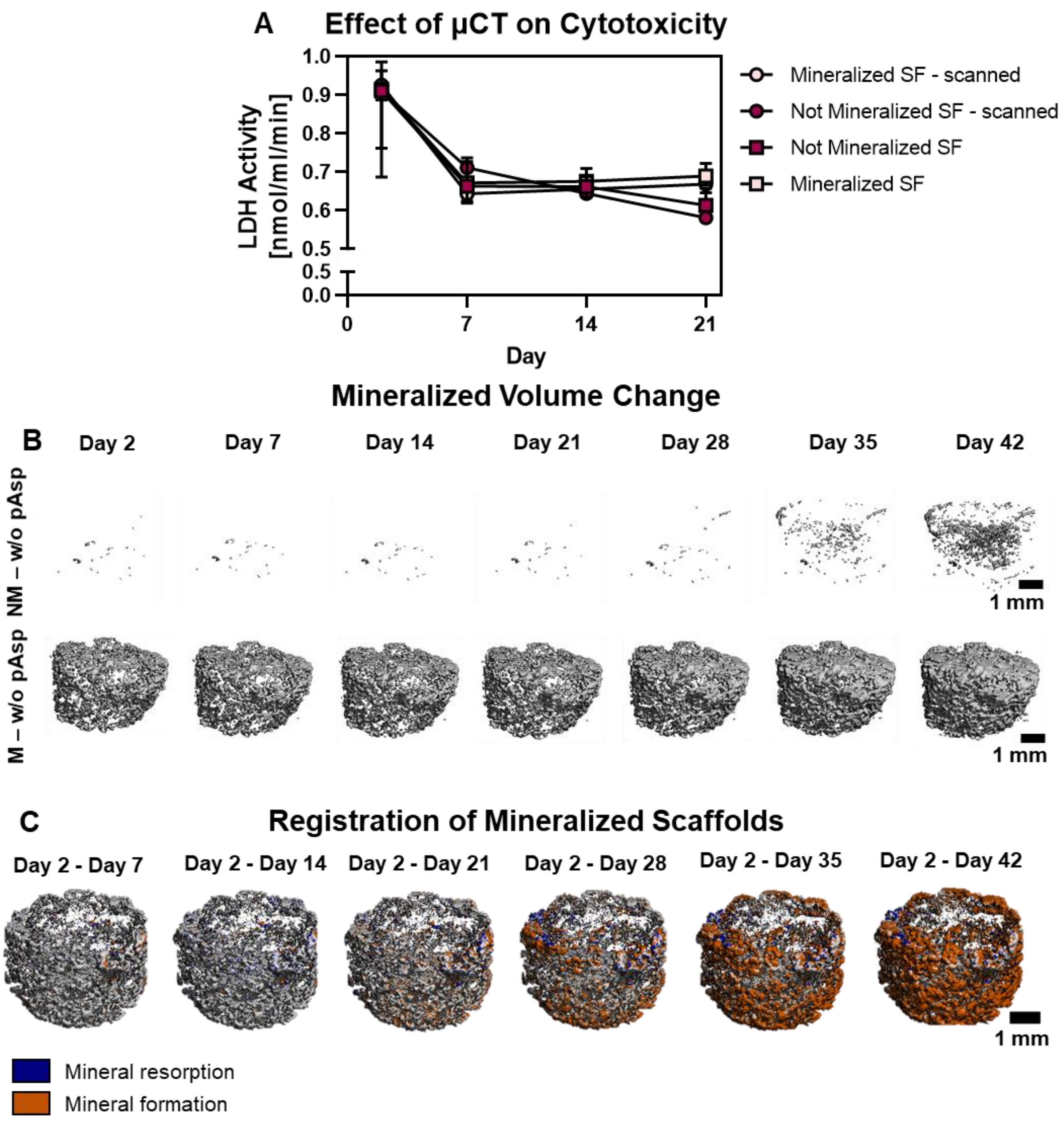


**Figure S6.** Longitudinal micro-computed tomography ($\mu$CT) scanning analyses. First, the influence of scanning on cell death was evaluated over 21 days (**A**). No differences between scanned and unscanned constructs were found, $\mu$CT scanning was therefore considered as a harmless method to track *in vitro* remodeling. Scans were segmented (**B**) and registered (**C**) to obtain information about mineralized volume change and resorption and formation sites, respectively.

A statistically significant higher sulphated glycosaminoglycan (GAG) content was found on day 21 of culture. GAGs were therefore visualized with an alcian blue staining. These GAGs were visualized between the trabecular-like structures (Figure S7).

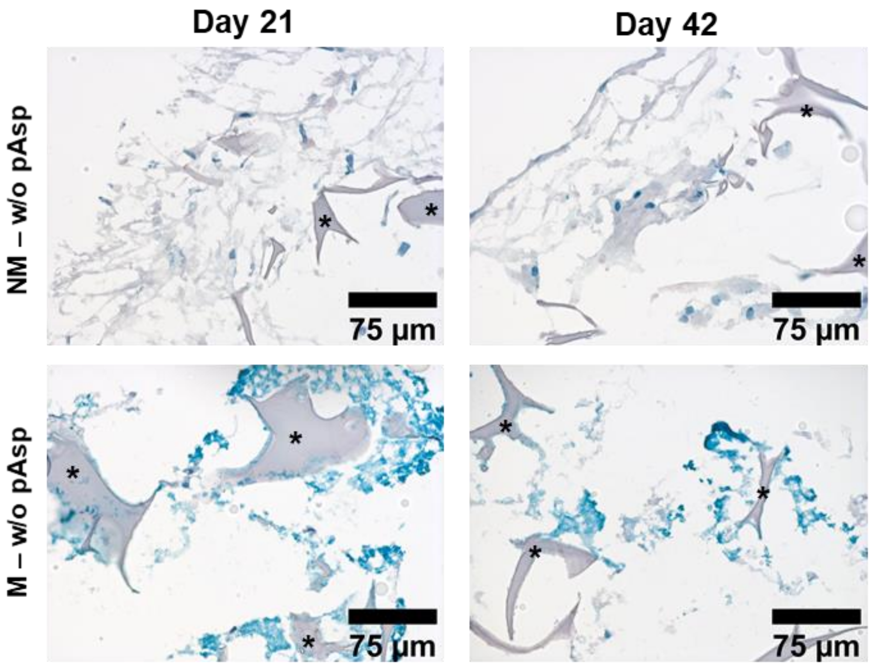


**Figure S7.** Glycosaminoglycan (GAG) visualization with alcian blue staining. GAGs were observed between the trabecular-like structures in mineralized (M) silk fibroin (SF) scaffolds on day 21 and 42. Asterisks indicate the scaffold trabeculae. Abbreviations: poly-aspartic acid (pAsp), non-mineralized (NM).

The antibodies that were used for immunofluorescent stainings are listed in Table S1.

**Table S1.** List of antibodies that were used in this study.

| Antigen | Supplier | Catalogue No. | Conjugate | Species | Dilution |
|---|---|---|---|---|---|
| RUNX2 | Abcam | ab23981 | | Rabbit | 1:500 |
| Osteopontin | Thermo Fisher | 14-9096-82 | | Mouse | 1:200 |
| Cathepsin K | Abcam | Ab37259 | | Mouse | 1:200 |
| Integrin-$\beta$3 | Abcam | Ab227702 | | Rabbit | 1:200 |
| Anti-mouse IgG1 | Molecular Probes | A21240 | Alexa 647 | Goat | 1:200 |
| Anti-Rabbit IgG | Molecular Probes | A21428 | Alexa 555 | Goat | 1:200 |
| Anti-mouse IgG2b | Molecular Probes | A21141 | Alexa 488 | Goat | 1:200 |

Abbreviations: runt-related transcription factor 2 (RUNX2)
